# Structural and functional insights into yeast Rqc1p, a protein required for thermotolerance with potential nuclear localization

**DOI:** 10.64898/2026.06.19.733457

**Authors:** Amanda Cristina Pereira-Antônio, Frederico Gabriel de Carvalho Oliveira, Marcelo Mesa Costa-Lima, Aysllan Fernandes Coelho, Erik Marques Rodrigues, Glória Regina Franco, Mario Henrique de Barros, Lucas Bleicher, Erich Birelli Tahara

## Abstract

Protein homeostasis – i.e., proteostasis – is the biological process by which the qualitative and quantitative balance of the proteome is conducted, either by preserving functionally relevant proteins or by degrading unnecessary ones. Stress conditions can modulate cellular proteostasis in order to promote cytoprotection and preserve the viability of living organisms. Among the cellular pathways already described that can play an important role in preserving biological functions by modulating proteostasis are the heat shock response and the ribosome quality control pathways. In this work, we show that the Rqc1p protein is necessary for the thermoadaptation of *S. cerevisiae* to heat shock, as *RQC1*-deficient yeast is sensitive to elevated temperatures. *In silico* approaches – such as multiple sequence alignment, structural analysis, and molecular dynamics simulations – confirmed earlier predictions that Rqc1p shares characteristics with the bHLH family of proteins. We also verified, through computational prediction of sub-cellular localization, that *S. cerevisiae* Rqc1p contains nuclear localization signals, suggesting that this protein can potentially be translocated toward the nucleus, thereby broadening its current range of recognized biological functions in this organism. Also, analysis of yeast transcriptomes subjected to heat shock showed that Rqc1p mRNA levels do not fluctuate in response to heat shock, suggesting that cellular concentrations of Rqc1p are already at optimal levels to elicit a rapid and effective response during thermal stress in *S. cerevisiae*.

## Introduction

One of the most fundamental characteristics of all life forms is their ability to adapt to the evolving conditions of their surroundings. These environmental fluctuations often trigger internal changes that disrupt the homeostasis of organisms, eliciting a wide range of vital responses, including remodeling of the cellular proteome (Gasch et al., 2000; Santiago et al., 2020). Different types of stress – such as thermal stress, oxidative stress, exposure to xenobiotics, and others – can modulate both the dynamic equilibrium and the regulation of specific protein groups within a cell. In fact, stressors alter cell proteostasis: proteins that become unnecessary in the new cellular context are subject to proteolysis, while those whose presence becomes indispensable – such as those harboring cytoprotective roles – are translated (Richter et al., 2010; Balchin et al., 2016).

Stress conditions frequently cause unfavorable modifications in protein structure, leading to the loss of their functional conformations. Therefore, to avoid potential cellular collapse due to extensive protein unfolding, cells exploit mechanisms that ensure protein refolding to prevent their proteome from being overtaken by defective proteins (Morimoto, 2008; Hartl et al., 2011). Heat shock proteins (HSPs) are the cellular agents responsible for this process as they play fundamental roles in protein folding and refolding, and also in preventing the formation of protein aggregates as well as in the disassembly of this cytotoxic species (Bukau et al., 2006; Yonashiro et al., 2016; Vihervaara et al., 2018; Alagar Boopathy et al., 2022).

A highly conserved pathway in living organisms called *heat shock response* (HSR) is responsible for the synthesis of several HSPs in situations of proteostatic collapse, such as those promoted by thermal, translational, chemical, genotoxic, and oxidative stress. In fact, the HSR pathway is required for a coordinated transcriptional response to the loss of protein homeostasis (Morimoto, 1998; Anckar & Sistonen, 2011; Gomez-Pastor et al., 2018). In eukaryotes, the HSR is mostly mediated by the transcription factor HSF1, which induces the expression of HSPs and other proteostasis genes in response to the accumulation of misfolded or aggregated proteins (Baler et al., 1992; Zheng et al., 2016). Interestingly, the HSR pathway is one component among several others in the cell proteostasis network, an interdependent system of cellular pathways that regulate the entire life cycle of proteins – from their synthesis to their degradation – and ensure their correct folding, proper subcellular localization, and elimination of damaged or potentially toxic species (Balch et al., 2008; Hartl et al., 2011; Sampaio-Marques & Ludovico, 2018).

Another member of the cell proteostasis network is the *ribosome quality control* (RQC) pathway, which detects newly-synthesized defective polypeptides stalled on ribosomes and polyubiquitinates them for proteasomal degradation (Bengtson & Joazeiro, 2010; Brandman et al., 2012). The translation of atypical mRNAs, such as those lacking stop codons – resulting mostly from premature polyadenylation events (Frischmeyer et al., 2002; van Hoof et al., 2002) – or those displaying tracts rich in rare codons, and also those harboring secondary/tertiary structure, causes polysomal collision and translational arrest, an event detected by Asc1p – a ribosomal protein belonging to the 40S subunit – cooperatively with the ribosome-associated ubiquitin E3 ligase Hel2p (Sitron et al., 2017; Joazeiro, 2019). Oligoubiquitination by Hel2p labels collided ribosomes, which become targets of the *ribosome quality control trigger* (RQT) complex, leading to dissociation of their two subunits (Matsuo et al., 2017; Winz et al., 2019). Afterwards, Rqc2p recognizes a stable peptidyl-tRNA bond at the P site of the 60S subunit (Lyumkis et al, 2014) and recruits the ubiquitin E3 ligase Ltn1p, which – in coordination with Rqc1p (Abaeva et al., 2025; Li et al., 2025) – promotes K48-linked polyubiquitination of the defective stalled polypeptides. Once marked for proteasomal degradation, these polypeptides are extracted by the AAA ATPase Cdc48p and its cofactors Ufd1p and Npl4p, which escort them to proteasome-mediated proteolysis (Defenouillère et al, 2013 and 2016; Tsuchiya et al., 2017; Kuroha et al., 2018; Joazeiro, 2019).

The RQC pathway becomes of pivotal importance in stress conditions due to the intense protein synthesis required to remodel the cellular proteome during these scenarios. In addition, it is known that various types of cellular stress increase the stabilization of mRNAs with structural folding, and also the generation of aberrant mRNAs – including those devoid of stop codons, or those truncated –, hindering translation elongation (Joazeiro, 2019; Monaghan et al., 2023). This increase in atypical mRNA synthesis results from direct defects in processing and transcription, as well as from the uncoupling between transcription, translation, and RNA surveillance, leading to robust activation of RNA and translation quality control pathways (Isken & Maquat, 2007; Decker & Parker, 2012; Inada, 2013).

To date, two known mechanisms interconnect the RQC pathway and the HSR: the first one is the accumulation of newly-synthesized defective polypeptides with CAT-tail, which, by stimulating the generation of protein aggregates, elicits proteotoxic stress and activates HSF1 (Defenouillère et al., 2013; Choe et al., 2016; Yonashiro et al., 2016); the second one is the unexpected Asc1p-mediated attenuation of Ssa4p synthesis – an Hsp70 – in situations of heat shock (Alagar Boopathy et al., 2023). However, the potential involvement of other factors from the RQC pathway in the HSR response remains unclear.

In this context, we sought to investigate whether any of the core elements of RQC complex play a role in yeast response to thermal stress. Surprisingly, we observed that Rqc1p is necessary to preserve the viability and replication ability of *S. cerevisiae* subjected to heat stress. We also verified the presence of a genetic interaction between *RQC1* and *LTN1,* as *rqc1*Δ*ltn1*Δ mutants rescued the thermoresistant, wild-type phenotype. Further *in silico* analysis indicated that Rqc1p harbors not only common structural traits of members from the bHLH family of proteins but also presents nuclear localization signals, suggesting that this protein may be translocated from the cytosol to the nucleus where it may play a role beyond its established functions in the cytosolic RQC pathway. Finally, the analysis of public yeast RNA sequencing data showed that *RQC1* transcription levels do not fluctuate when *S. cerevisiae* wild-type is exposed to high temperatures, suggesting that Rqc1p is already present at an optimal cellular concentration to enable a rapid and efficient response to heat shock in this organism.

## Materials and Methods

### Strains and growth media

The *Escherichia coli* strain used for molecular cloning was RR1 (Bolivar et al., 1977). The genotypes of yeast strains used in this study are provided in Table S1. *Saccharomyces cerevisiae* BY4741 strains harboring single deletions were kindly donated by Prof. Luis Eduardo Soares Netto from Instituto de Biociências – Universidade de São Paulo (São Paulo, SP, Brazil), and the double deleted-gene mutants were previously generated by our group (Valdez, 2018). The culture media used for *E. coli* were LB (1% tryptone, 1% NaCl, and 0.5% yeast extract), and LB supplemented with ampicillin (100 μg/mL). Culture media used for *S. cerevisiae* were YPD (1% yeast extract, 2% bacteriological peptone, and 2% dextrose), and SD-Leu (0,67% yeast nitrogen base with ammonium sulfate, 2% dextrose, 10 mg/L adenine sulfate, 20 mg/L uracil, 50 mg/L tryptophan, 20 mg/L histidine, 50 mg/L arginine, 50 mg/L tyrosine, 50 mg/L lysine, 50 mg/L phenylalanine, 1 g/L glutamate, 1 g/L aspartate, 140 mg/L valine, 100 mg/L threonine, 4 g/L serine, 20 mg/L methionine, 50 mg/L isoleucine, and 80 mg/L asparagine).

### Spot assays and maximum specific growth rate determination

Yeast tolerance to heat stress was assessed both by spot assays (Kwolek-Mirek & Zadrag-Tekza, 2014) on solid media and by quantifying the maximum specific growth rate (m^max^/h) in liquid cultures (Monod, 1949). Spot assays were carried out by plating 3 or 5 mL of serially-diluted cells suspensions onto solid YPD or SD-Leu using cells grown overnight (16 h) in respective liquid media at 28°C.

Photographic documentation was performed after 48 h of growth at 28°C (control) or 38°C (heat stress). To determine μ^max^/h, yeast cells were pre-grown in liquid YPD for 16 h and re-inoculated in fresh medium with initial Abs_600nm_ = 0.0025. After 12 h of culture either in control or in heat shock conditions, the values of Abs_600nm_ were assessed every hour until 24 h of growth, and were then converted to their natural logarithm (ln). Growth curves were generated by plotting the Abs_600nm_ values (in ln) in the ordinate and the time values (in h) in the abscissa, and its respective μ^max^/h was resolved through the determination of the slope of the linear regression obtained using the points belonging to the linear segment of the growth curve.

### Multiple sequence alignment and secondary structure prediction

The protein sequences from the TCF25 family were obtained from The UniProt Consortium (https://www.uniprot.org). “TCF25”, “RQC1”, and “NULP1” were the keywords used in the search, alongside the names of eukaryotic model organisms to represent biological diversity. Selected sequences are listed in Table S2. The multiple sequence alignment was performed using MUSCLE (Edgar, 2004; Madeira et al., 2024), and the results were processed and visualized using Jalview (Waterhouse et al., 2009). NetSurfP-3.0 (Høie et al., 2022) was used to predict the secondary structure and *relative surface accessibility* (RSA) of Rqc1p.

### Molecular cloning

The *RQC1* gene sequence was obtained from SGD (*Saccharomyces* Genome Database) (Cherry et al., 1998). The sequence and the utilization of each primer used in this study are indicated in Table S3. *S. cerevisiae* genomic DNA was obtained as described by Harju et al. (2004). Polymerase chain reactions were performed using either Platinum^TM^ Taq Polymerase (Invitrogen) or Phusion High-Fidelity DNA Polymerase (New England Biolabs), according to the manufacturers’ protocols. Amplicons were purified by electroelution from an 1% agarose gel, followed by phenol-chloroform DNA extraction. Restriction enzyme digestions of amplicons and plasmid DNA were carried out using HindIII, PstI, and KpnI (New England Biolabs), or EcoRI (Thermo Fisher Scientific), and plasmid digestions were performed in the presence of alkaline phosphatase (Roche). Ligation of DNA fragments was conducted using a recombinant DNA ligase produced in-house in the presence of a commercially available ligase buffer (New England Biolabs). Competent *E. coli* was prepared and transformed as described by Hanaham (1983), and positive clones were confirmed by diagnostic restriction digestion analysis and Sanger DNA sequencing (Genoma USP, São Paulo, SP, Brazil), using plasmids extracted with the PureYield^TM^ Plasmid Miniprep System (Promega) from selected *E. coli* clones. *S. cerevisiae* transformation was conducted according to Gietz and Schiestl (2007).

### Residue conservation and coevolution analyses

A multiple sequence alignment of the Tcf25 domain from various organisms was generated by querying UniProt (The UniProt Consortium, 2025) for the Pfam domain PF04910 and aligning the resulting sequences applying the Hidden Markov Model through HMMER (Eddy, 2011). This alignment served as the basis for conservation and coevolution analyses conducted with PFstats (Fonseca-Júnior et al., 2018) using the following protocol: sequences with less than 80% of identity to Tcf25 or more than 80% of identity to other sequences in the alignment were removed to eliminate fragments and redundancy. A Dima-Thirumalai test (Dima & Thirumalai, 2006) established that a minimum sub-alignment size of 15% was statistically significant, which was then applied in the *Decomposition of Residue Coevolution Network* (DRCN) analysis (Bleicher et al., 2011). Residue pairs were considered as coevolving when (i) both residues appeared in at least 15% of the sequences; (ii) the presence of each residue increased the frequency of the other to at least 80%; and (iii) the associated p-value was below 10^-10^, simultaneously. A coevolutionary network was constructed with residue–position pairs as nodes which were connected when they exhibited statistically significant coevolution according to the aforementioned thresholds. The networks were decomposed into connected components, each representing groups of specific residues at defined positions that co-occur across subsets of proteins in the alignment. These modules correspond to sets of coevolving residues, typically associated with functional or structural roles.

### Three-dimensional structure analysis

A recently published 3.16Å structure of the entire RQC complex using cryo-electron microscopy (PDB code: 9OFV) by Li et al. (2025) was employed for *S. cerevisiae* Rqc1p structural analysis using PyMOL (Schrödinger, LLC).

### Sequence analysis, homology modeling and molecular dynamics simulations

The amino acid sequence of Rqc1p was aligned against proteins harboring helix-loop-helix (bHLH) domains available on Protein Data Bank (PDB) as of May 2026 through pairwise sequence alignment. The region comprised by residues Glu127-Ile217 exhibited similarity to the bHLH domain of the human microphthalmia-associated transcription factor (MITF; PDB: 7D8T), although with a very low score (-10), determined by BLOSUM62 substitution matrix with gap opening and extension penalties of −15 and −2, respectively. To assess the statistical significance of the alignment, one of the aligned sequences was randomly shuffled and realigned to its counterpart, and the best alignment score was recorded. This procedure was repeated 10^6^ times to generate a null distribution of alignment scores. As negative controls, segments derived from the N- and C-terminal regions of Rqc1p with equivalent lengths (i.e., containing 91 residues) were subjected to the same shuffling and alignment protocol. The putative bHLH-containing region of Rqc1p was subsequently subjected to comparative structural modeling through MODELLER (Šali & Blundell, 1993), using the MITF structure as template. Three independent modeling systems were generated: (i) a putative Rqc1p bHLH homodimer comprising residues 123–217; and (ii–iii) negative-control models generated from N- and C-terminal Rqc1p segments threaded onto the same MITF template. For each modeling system, 10^3^ structural models were generated, and the lowest-energy ones were selected for subsequent analyses. Selected models were parameterized using the CHARMM36 force field (Best et al., 2012) through the CHARMM-GUI (Jo et al., 2008) interface and solvated in TIP3P water boxes. Energy minimization was performed in order to remove steric clashes and optimize local geometry. Systems were then equilibrated for 1 ns under NVT conditions, followed by 100 ns production simulations under NPT conditions at 303.15 K and 1 bar pressure. All molecular dynamics simulations were performed using NAMD (Philips et al., 2020).

### Subcellular localization prediction

DeepLoc 2.1 (Ødum et al., 2024) and NucPred (Brameier et al., 2007) were used to predict subcellular localization and the presence of sorting signals in Rqc1p, while potential NLSs were predicted using cNLS Mapper (Kosugi et al., 2009).

### Transcriptome analyses

RNA sequencing data from wild-type *S. cerevisiae* challenged with heat shock were retrieved from the *sequence read archive* (SRA) using the MeSH terms “*Saccharomyces cerevisiae*” AND “heat shock”. To ensure greater homogeneity and robustness in the analyses, a manual curation was performed to identify only studies that used paired-end sequencing and had at least three biological replicates per condition. The quality of the RNA-seq libraries was analyzed using FastQC v.0.11.9 (https://www.bioinformatics.babraham.ac.uk/projects/fastqc/) and the MultiQC v.1.13.dev0 (Ewels et al., 2016). After that, the total of four studies were selected and are summarized in Table S4. RNA-seq library read mappings were performed against the *S. cerevisiae* reference genome of the S288C strain (NCBI entry code: GCF_000146045.2) using STAR v.2.7.10b (Dobin et al., 2013). Conversion and sorting of BAM files to SAM were performed using Samtools v.1.16.1 (Danecek et al., 2021). The counting of the number of mapped reads per gene was performed using FeatureCounts v.2.0.3 (Liao et al., 2014). The resulting count matrix was imported into R environment for downstream analyses. Investigation of differential gene expression was performed using the DESeq2 v.1.40.2 package (Love et al., 2014). To consider a gene as differentially expressed, we adopted a cutoff of log_2_ (FoldChange) ≥ 1, and an adjusted p-value (padj) < 0.05.

## Results

### Rqc1p is required for *S. cerevisiae* thermotolerance

We first sought to investigate whether the integrity of the RQC complex is necessary for *S. cerevisiae* thermotolerance to heat stress. To do so, we used cells lacking *RQC2*, *LTN1*, and *RQC1* to perform a heat stress tolerance assessment by conducting a spot assay in which these mutants, along with the wild-type cells, were spotted onto YPD plates and incubated either at 28°C (standard growth temperature) or at 38°C (heat stress temperature). After 48 h, we observed that the growth of wild-type, *rqc2*Δ, and *ltn1*Δ cells was comparable at both incubation temperatures, whereas the *rqc1*Δ mutant showed reduced growth at 38°C compared with 28°C. These results indicate that Rqc1p is required for *S. cerevisiae* response to heat stress and is thus involved in yeast cytoprotection upon supraoptimal temperature growth conditions (Figure 1A).

**Figure 1.**
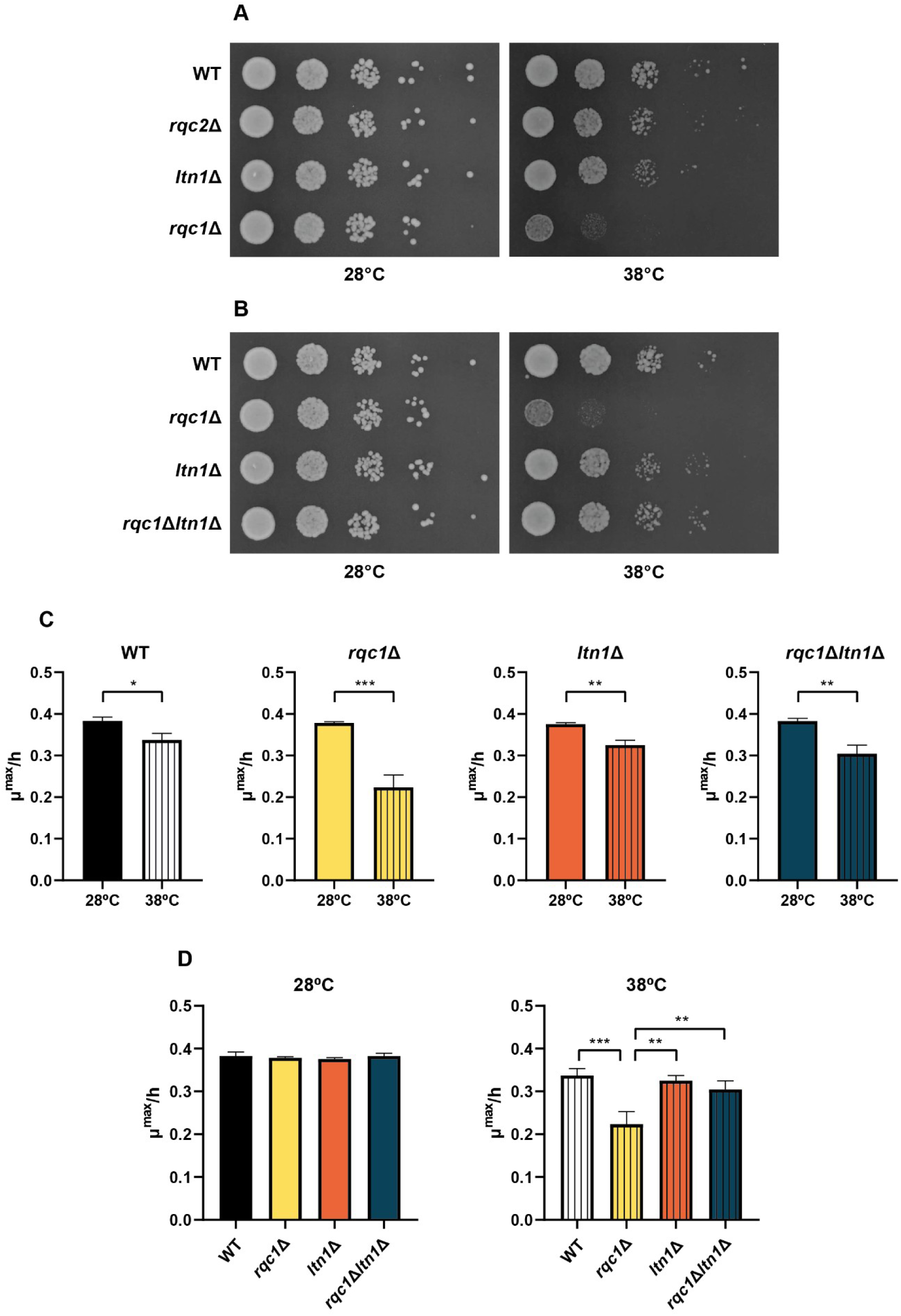
Rqc1p is required for *S. cerevisiae* thermotolerance. **Panels A-B:** Spot assay of RQC complex mutants at 28°C and 38°C (heat stress) on rich glucose medium (YPD). Plates were photographed after 48 h. **Panel C:** Comparative analysis of the maximum specific growth rate (μ^max^/h) for WT, *rqc1*Δ, *ltn1*Δ, and *rqc1*Δ*ltn1*Δ cells at 28°C and 38°C (unpaired Student’s *t*-test). **Panel D:** Comparison of μ^max^/h among all strains at 28°C and 38°C (one-way ANOVA/Tukey). Statistical significance was defined as: *p < 0.05, **p < 0.01, and ***p < 0.001.

Rqc1p was first implicated in the RQC complex via a genome-wide screen, which revealed that *ltn1*Δ and *rqc1*Δ mutants share a highly analogous interaction profile (Brandman et al., 2012); we then hypothesized that the deletion of *RQC1* might provide a combined effect on the heat shock response in *LTN1*-deficient cells. In fact, it is currently known that Ltn1p-generated K48-type polyubiquitin chains – those which are identified as targets by Cdc48p and its cofactors Ufd1p and Npl4p for extraction of the halted translational complex – are preferentially formed in the presence of Rqc1p (Abaeva et al., 2025), indicating that these two proteins act in a coordinated fashion. Therefore, we constructed the double mutant *rqc1*Δ*ltn1*Δ to investigate the possible effects of *LTN1* deletion on the resistance of *RQC1*-deficient cells when grown in supraoptimal temperature conditions.

Surprisingly, the double mutant *rqc1*Δ*ltn1*Δ demonstrated increased resistance when cultivated at 38°C when compared to the single mutant *rqc1*Δ, displaying comparable growth to that of wild-type and *ltn1*Δ cells, which indicates the existence of a genetic interaction between *LTN1* and *RQC1* (Figure 1B). This positive genetic interaction suggests that inhibiting polyubiquitination in *RQC1*-deficient yeast confers increased thermotolerance, although the underlying mechanisms responsible by this phenotype remain to be elucidated. Nevertheless, the suppression of polyubiquitinated aggregate accumulation in *rqc1*Δ cells by *LTN1* deletion previously observed (Defenouillère et al., 2016) is recapitulated here, further supporting an epistatic relationship between these two genes.

To quantitatively confirm the semiquantitative spot assay results, and upon the understanding that greater sensitivity to thermal shock would result in a reduced growth rate, we determined the maximum specific growth rate (μ^max^/h) of wild-type, *rqc1*Δ, *ltn1*Δ, and *rqc1*Δ*ltn1*Δ cells cultured both at 28°C and 38°C in liquid YPD under constant orbital agitation. We observed that, when cultured at 38°C, all cells showed a significant decrease in μ^max^/h compared with those at 28°C (Figure 1C). In addition, we verified that the *rqc1*Δ mutants presented, in fact, a significant reduction in their m^max^/h when compared to those determined for wild-type, *ltn1*Δ, and *rqc1*Δ*ltn1*Δ cells when cultured upon supraoptimal temperature conditions (Figure 1D). Altogether, these data corroborate our previous observation that Rqc1p is critical for *S. cerevisiae* to properly cope with heat stress.

### *S. cerevisiae* Rqc1p exhibits conserved sequence features and a bipartite domain organization

During the 2000s, orthologs of Rqc1p – first annotated as Nuclear Localized Protein 1, or NULP1, and later standardized in genomic databases as TCF25 – had been identified in mice and humans, and described as transcription factors belonging to the basic helix-loop-helix (bHLH) family (Olsson et al., 2002; Cai et al., 2006). Members of the bHLH family contain a tract of basic amino acid residues – Lys and Arg – which is usually responsible for their DNA binding, followed by a helix-turn-helix structural motif which mediates protein dimerization (Staudinger et al., 1993; Robinson & Lopes, 2000; Jones, 2004). These features are associated with the regulation of gene transcription during cellular differentiation and development by these proteins (Murre et al., 1994 and 2019). Notably, the *S. cerevisiae* Rqc1p was included in those foundational comparative analyses on the bHLH family, suggesting the conservation of sequence features across distant species (Cai et al., 2006, Zeng et al., 2017). We therefore revisited these early annotations using multiple sequence alignment of Rqc1p/TCF25 homologs across representative species using MUSCLE (Edgar, 2004; Madeira et al., 2024) aiming to identify conserved residues and domains shared across orthologs in selected model organisms. This analysis was conducted to (i) investigate evolutionary conservation, (ii) further support orthology across eukaryotes, and (iii) identify potentially functional regions in Rqc1p.

We then identified the existence of two highly conserved regions in *S. cerevisiae* Rqc1p: the first one, located near to its N-terminal – spanning residues 92 to 107 – characterized by the presence of multiple Lys and Arg residues, thus constituting a polybasic tract (Figure 2A); and the second one, located downstream – spanning residues 328 to 663 –, enriched in hydrophobic and polar amino acid residues (Figure 2B). Interestingly, this latter conserved region of Rqc1p exhibits strong sequence similarity to the Tcf25 domain (PF04910), as defined by the Pfam database, indicating conservation of a structural module present in Rqc1p/TCF25 orthologs.

**Figure 2.**
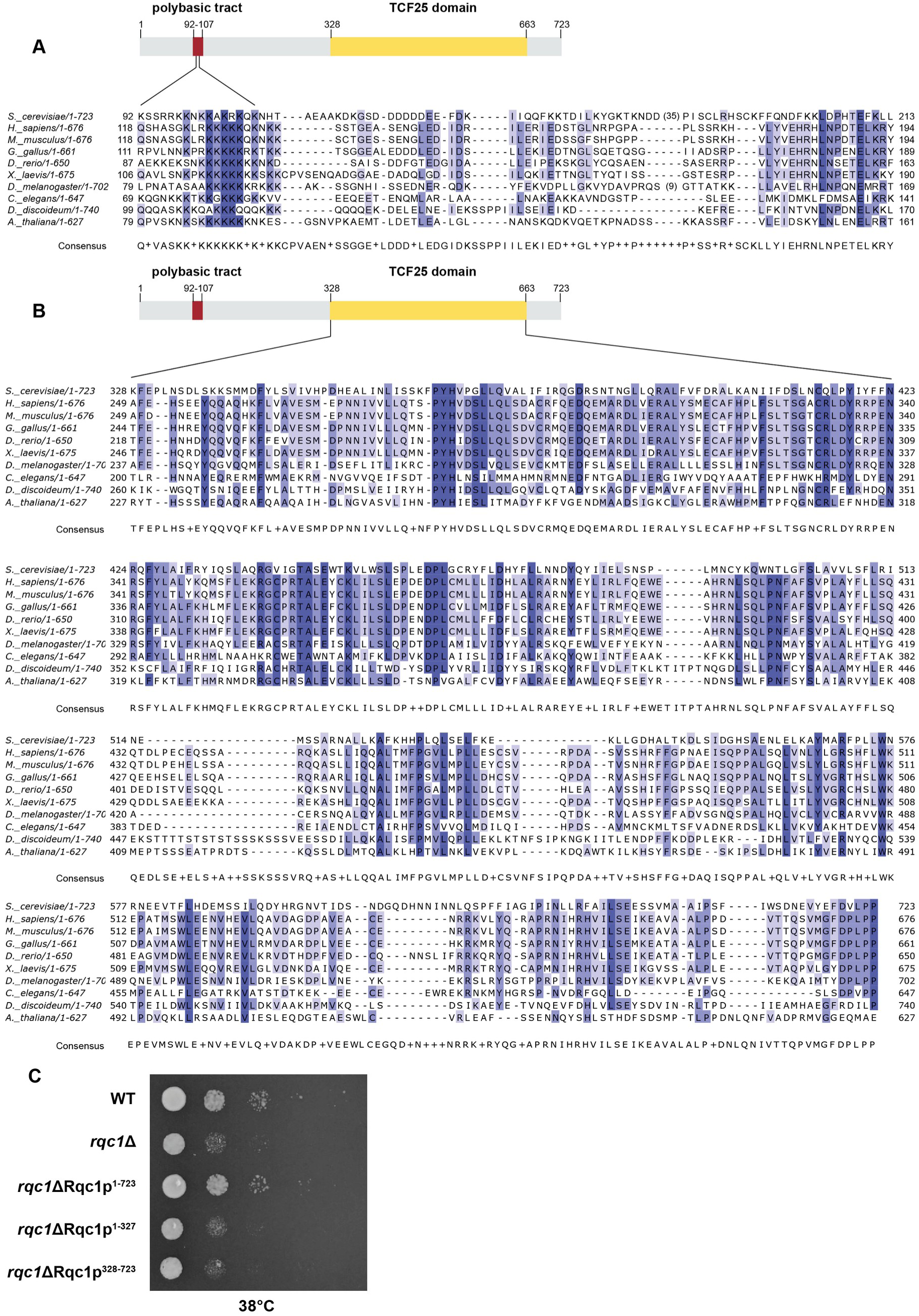
Rqc1p exhibits conserved sequence features and a bipartite domain organization, with both regions being required for thermotolerance. **Panels A-B:** Schematic representation of domain composition and multiple sequence alignment of Rqc1 homologues, highlighting the polybasic tract (A) and the Tcf25 domain (B). Percentage of identity is indicated (blue gradient). **Panel C:** Spot assay of Rqc1p mutants at 38°C (heat stress) on auxotrophic medium (SD-Leu). Both the wild-type (WT) cells and the *rqc1*Δ mutants were transformed with an empty vector (YCplac111) harboring only the promoter and terminator regions, without the complete *RQC1* coding sequence. Transformants obtained using YCplac111 constructs encoding either the Rqc1p full-length (1-723) or partial (1-327 and 328-723) coding regions are indicated in the figure. Plates were photographed after 48 h.

To investigate whether each of these two highly conserved regions may contribute independently to Rqc1p-mediated thermotolerance in yeast, we tested if the expression of a single Rqc1p region could prevent the increased sensitivity of *RQC1*-deficient cells to thermal stress. To do so, we transformed *rqc1*Δ mutants with specific centromeric constructs encoding (i) either the full-length Rqc1p; or (ii) the N-terminal region harboring the Arg/Lys polybasic tract; or (iii) its Tcf25 domain; and conducted another spot assay in which these yeast transformants were spotted onto auxotrophic SD plates and incubated at 38°C for 48 h. We observed that, while the construct harboring the full-length *RQC1* restored heat stress tolerance in *rqc1*Δ cells, the constructs encoding the N-terminal region or the Tcf25 domain did not (Figure 2C). These results therefore suggest that neither isolated Rqc1p region is able to rescue the wild-type phenotype in *S. cerevisiae* cells devoid of *RQC1* under heat stress, suggesting that thermotolerance in this yeast requires the coordinated action of both conserved Rqc1p regions.

### Coevolutionary analysis of *RQC1* homologs revealed nine communities of coevolving residues and three highly-conserved residues

Neofunctionalization often results from gene duplication followed by gene divergence (Hughes 1994; Walsh, 2003). After querying OrthoDB (Kusnetsov et al., 2023) for *RQC1* homologs, we found that their coding genes are single-copy in most species: 2450 out of 2538 species cataloged on OrthoDB harbor a single TCF25 homolog, suggesting that duplication events within this gene family were infrequent during evolution. Thus, novel biological roles could emerge only from mutations in specific residues that did not disrupt preexisting functional domains, since most species lack a second gene copy to retain the ancestral function. In these situations, residue coevolution analysis could detect such events, as mutations in functional sites would yield coevolutionary signals (Marks et al., 2012; de Juan et al., 2013). We next performed a residue coevolution analysis of TCF25 homologs retrieved from OrthoDB to identify mutations at critical sites that may be compensated by substitutions at other residues, thereby enabling structural adaptations while preserving protein function using DRCN (Bleicher et al., 2011). Although we were unable to analyze both termini ends of TCF25 as they exhibit low sequence conservation, we successfully investigated coevolutionary patterns within the Tcf25 domain. The results revealed nine distinct communities of coevolving residues within this region (Figure S1). A community of coevolving residues – or a coevolutionary community – is a group of residues that are densely interconnected with one another than with the rest of the protein network (Newman & Girvan, 2004; da Fonseca et al., 2019).

Notably, we found that residues comprising the coevolutionary community 1 are highly conserved across the analyzed species, with this conservation pattern being largely maintained from yeasts to humans (Figure 3A). Through the examination of the cryo-EM structure of Rqc1p within the RQC complex recently published by Li et al. (2025), we identified a direct functional role for one of the identified coevolved residues from community 1: Gln374 forms a hydrogen bond with Thr9 from ubiquitin (Figure 3B), suggesting that this Rqc1p residue contributes to protein-protein stabilization within the RQC complex.

**Figure 3.**
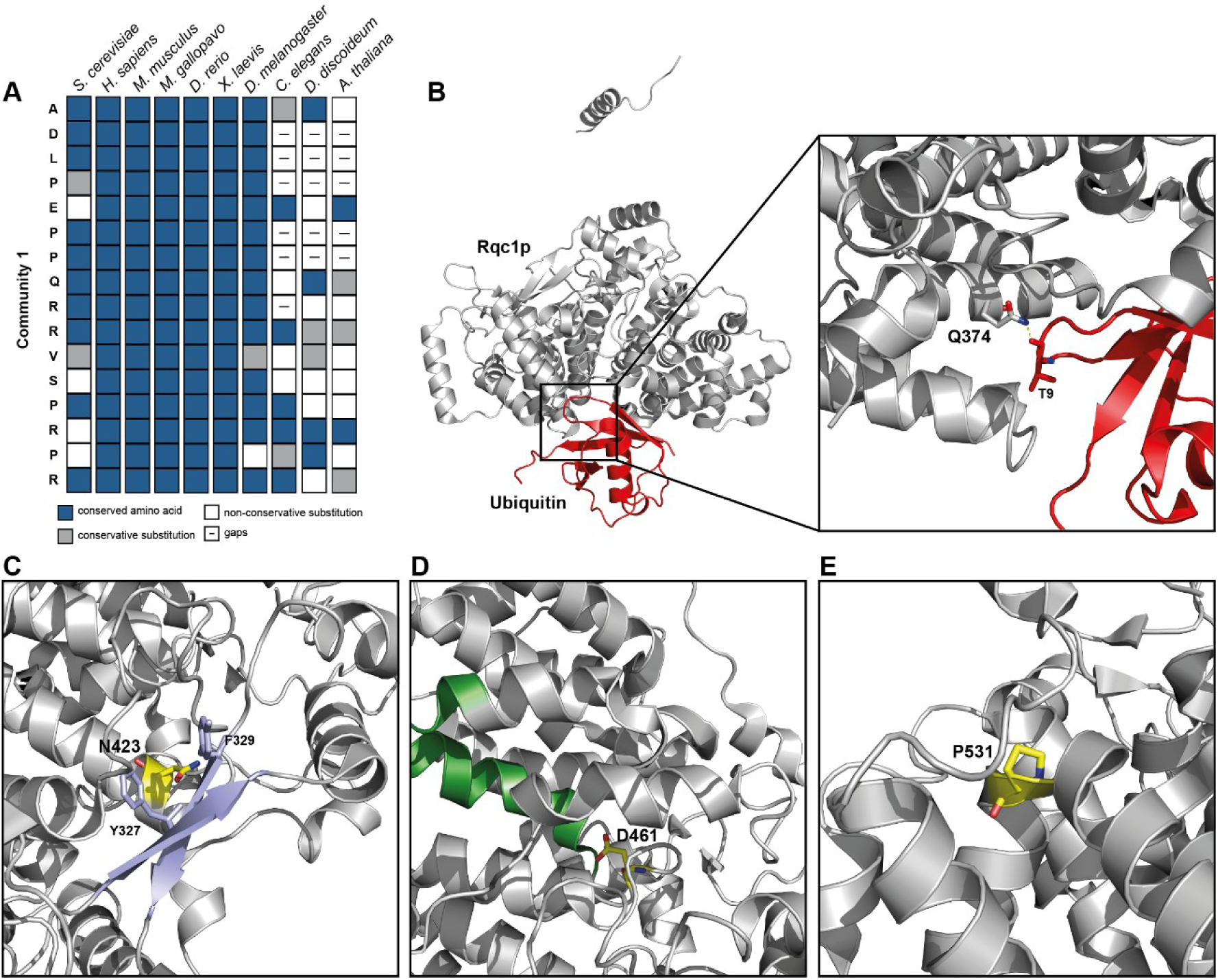
Conserved and coevolving residues stabilize key structural elements of the Tcf25 domain and mediate interaction with ubiquitin. Structural analyses were performed using the cryo-EM map of Rqc1p within the RQC complex (9OFV) available from Li et al. (2025). **Panel A:** Conservation pattern of residues belonging to coevolutionary community 1 across representative species. Boxes indicate conserved amino acids (blue), conservative substitutions (gray), non-conservative substitutions (white), and gaps (dashed). **Panel B:** The coevolved residue Q374 of community 1 within the Rqc1p (gray) forms a hydrogen bond with T9 of ubiquitin (red). **Panel C:** Conserved residue N423 (yellow) interacts with Y327 and F329 within a β-strand region, contributing to stabilization of the sole β-sheet element (light blue) identified in the structure. **Panel D:** Conserved residue D461 (yellow) stabilizes the helix (green) positive dipole. **Panel E:** Conserved residue P531 (yellow) introduces a structural kink between adjacent helical segments, potentially contributing to local structural organization.

In addition, since the DRCN pipeline allows the quantification of residue conservation across analyzed homologs, we further identified three conserved residues with potential importance: (i) Asn 423; (ii) Asp461; and (iii) Pro531, which are conserved in 83%, 95% and 81% of Rqc1p homologs examined, respectively. We then found that Asn423 interacts with Tyr327 and Phe329 within the beta-strand spanning residues 327-331, which in turn associates with another beta-strand formed by residues 287-291 to generate the sole protein segment organized in beta-sheet (Figure 3C). Moreover, we observed that Asp461 stabilizes the positive dipole at the N-terminus of a helix initiated at Leu464 (Figure 3D). Interestingly, despite their polar nature, both Asn423 and Asp461 are buried within the Rqc1p core, suggesting their role as relevant residues for structural stabilization of the protein. Finally, we also found that Pro531 creates a structural kink between adjacent helical segments (Figure 3E). Together, these findings suggest that the conserved residues in the Tcf25 domain contributed to maintaining the structure and functional properties of critical Rqc1p conformational elements, including β-sheet organization, helix stabilization, and conformational flexibility during evolution.

### Structural features of Rqc1p reveal potential determinants of RQC complex assembly and protein function

To further identify relevant insights into the molecular basis of Rqc1p, we revisited the cryo-EM structure of the RQC complex (Li et al., 2025) previously analyzed to identify the structural and functional roles of the co-evolved and conserved residues (Figure 3B-E). Interestingly, we then found that the extent of the Tcf25 domain of Rqc1p in the experimentally determined structure – i.e., the tract comprised between residues 176 to 682 – goes beyond the previously described predicted limits according to the Pfam database (residues 328 to 663). In fact, data available from the cryo-EM map indicates that the segments between residues 176 to 326 and 664 to 682 of Rqc1p are potential constituents of the Tcf25 domain as both of them participate to the same folding unit since, in the structure spanning from residue 176 to 682, flexible linkers are absent, and structural continuity, as well as topologic interdependence, are existent – this is consistent with the presence of a cooperatively folded structural unit between residues 176 and 682 (Figure 4A-B). In addition, we found that several residues located upstream of the previously annotated Tcf25 domain (residues 176 to 326) contribute to the interaction of Rqc1p with its RQC complex partners (Figure 4C): its Arg249 interacts through a hydrogen bond with the Cit2 from JA/f chain of 25S rRNA; its Lys254 establishes a hydrogen bond with Gln34 of the ribosomal protein L19-A; the carbonyl group of its Leu213 interacts with Lys1525 from Ltn1p via a hydrogen bond; its Asp215 is involved with Ser1524 and Lys1525 of Ltn1p through a hydrogen bond and a salt bridge, respectively. In addition, Asp216, Asp224, and Asp226 of Rqc1p establish salt bridges with Lys1521, Lys1541, and Arg1544 of Ltn1p, respectively; and Ser218 and Ser221 of Rqc1p interact with Asp1519 of Ltn1p through hydrogen bonds. Finally, we also observed that Ser639, Val655, Glu657, and Gln374 form hydrogen bonds with Thr66, Thr12, Gly10, and Thr9 from ubiquitin, respectively.

**Figure 4.**
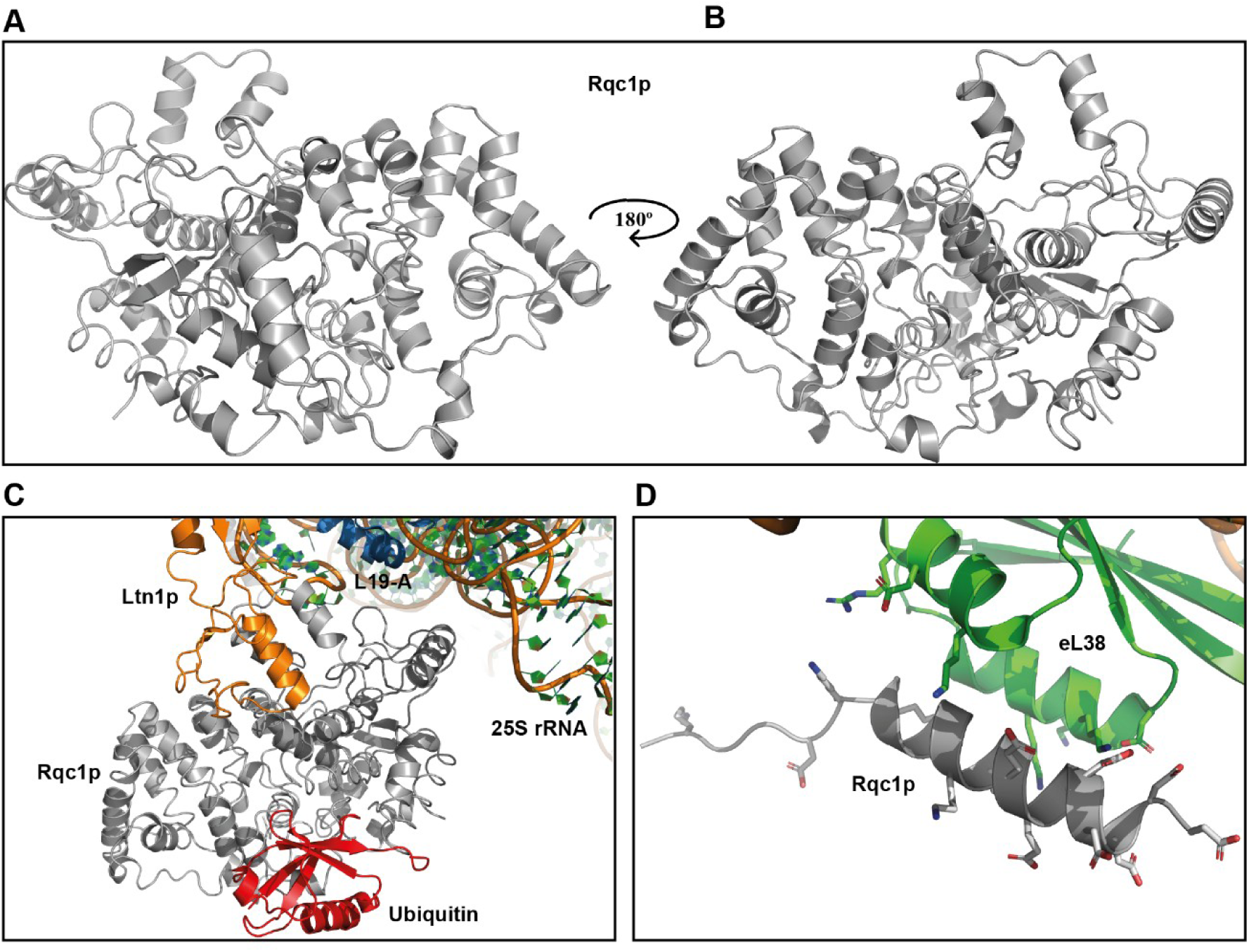
Rqc1p experimental structure in the context of the ribosome quality control context. **Panel A:** Three-dimensional structure of the 176-682 segment of Rqc1p. **Panel B:** Same structure as in Panel A after a 180° rotation. **Panel C:** Rqc1p (grey) binding partners on the ribosome quality control complex. The polynucleotide chain corresponds to 25S rRNA, and the colored protein chains correspond to ribosomal protein L19-A (blue), Ltn1p (orange) and ubiquitin (red). **Panel D:** The Rqc1p polyacidic 120-143 stretch (grey) as interpreted from the cryo-EM map establishing contacts with basic residues from eL38 (green). Charged residues for both proteins are shown as sticks for the sake of clarity.

As for the N-terminal domain of Rqc1p, we already had observed the presence of a non-conserved, enriched segment containing eight acidic residues between positions 120 and 127 in the previously performed multiple sequence alignment (Figure 2A). In addition, analyzing the cryo-EM map of the RQC complex, we also verified that this particular region of Rqc1p was assembled based on the cryo-EM density of ribosomal protein eL38, which harbors several basic residues that are in direct contact with the acidic ones from Rqc1p found in the aforementioned segment (Figure 4D). We then hypothesized that this region may undergo folding only when Rqc1p is associated with the other components of the RQC complex – particularly with eL38 –, since it is expected that acidic-rich segments – or basic-rich ones – are structurally organized as coils/random coils as a result of the presence of multiple neighboring residues with the same net charge when proteins are not interacting with partners (Creighton, 1993). To confirm this hypothesis, we performed a secondary structure prediction of unbound Rqc1p using NetSurfP-3.0 (Høie et al., 2022) to investigate the degree of solvent exposure and the structural disorder of this particular region of the protein.

The result of this analysis indicates that the residues found both in the N- and C-terminal ends of Rqc1p – which includes the 120-127 polyacidic stretch –, show high relative surface accessibility (RSA) values (> 0.25) and are thus favorably exposed to the solvent (Figure S2). Protein segments harboring such a characteristic frequently exhibit high conformational flexibility when not associated with their molecular partners; interestingly, the Rqc1p segment comprised between residues 1-119, which is immediately upstream to the 120-127 tract – i.e., its very N-terminal end –, does not show any visible density in the cryo-EM map, indicating that the 119 first residues of Rqc1p do not establish stable interactions within the resolved RQC complex structure, suggesting that its polybasic region (enriched with Lys/Arg) – which is located between residues 92 and 107 (Figure 2A) – may not be involved in the RQC complex assembly. Notably, Lys/Arg polybasic stretches are often used to promote electrostatic interactions between proteins and nucleic acids, being commonly observed in DNA-binding domains, nuclear basic peptides and nuclear localization signals (Pabo and Sauer, 1992; Luscombe et al., 2000).

### A putative HLH-like region in Rqc1p displays structural stability during molecular dynamics simulations

As previously mentioned, the earliest comparative analyses of TCF25 included *S. cerevisiae* Rqc1p as a member of the bHLH family; however, we verified that the cryo-EM map of the RQC complex shows weak to absent density for the Rqc1p N-terminal end, where its HLH domain is presumably located. Then, we evaluated whether unbound yeast Rqc1p may adopt a canonical HLH fold by simulating its structural stability through a molecular dynamics simulation.

First, we sought to identify regions within the primary structure of Rqc1p exhibiting similarity to HLH-containing proteins deposited in the PDB through sequence alignment analysis. The Rqc1p segment between residues Glu127 and Ile217 showed the highest propensity to align to an HLH structure found in the microphthalmia-associated transcription factor (MITF), a protein from the bHLH-Zip family (Hodgkinson et al., 1993). This alignment showed a score of -10, considering the BLOSUM62 matrix, and -15/-2 penalties for gap openings and extensions, respectively. Since this negative score suggests the existence of a potentially random similarity, we verified whether this alignment was biologically significant or occurred by chance. Thus, one of the two compared sequences was randomly shuffled and the best alignment was recalculated, with this entire process being repeated a total of 10 ^6^ times, in order to obtain alignment score values that occur purely by chance. The analysis showed that the actual alignment score is 4.02 standard deviations higher than the average of score values (Figure 5A). This suggests that the original alignment is statistically significant and did not occur by chance, and also that both protein segments have an evolutionary relationship (i.e., homology) or a strong pressure to maintain this specific conformation (i.e., functional convergence). As negative controls, the same procedure was applied to equally-sized segments – harboring 91 residues – from the N- and C-terminal regions. Not only the segments from these two regions presented much lower alignment scores with MITF (-42 for N-, and -55 for the C-terminal) but also scores that are similar to those obtained by shuffling (Figures S3A and S3C). This indicates that other Rqc1p regions would not yield a significant alignment such as that observed for the Rqc1p segment spanning residues 127-217. We next performed molecular modeling of Rqc1p residues 127-217 using MODELLER (Šali & Blundell, 1993), using as a template the structure of MITF linked to its responsive element, obtained from the PDB under the code PDB 7D8T (ID), as well as the two terminals previously used as negative controls. We observed that the Glu127-Ile217 segment of Rqc1p remained structurally stable and generally preserved its HLH arrangement along the trajectory in the absence of binding partners (Figure 5B). In contrast, both the negative controls did not maintain the HLH structure observed in the MITF template, being subjected to progressive structural disorganization during the simulation (Figures S3B and S3D). Taken together, these results provide further support for the existence of a putative HLH-like conformation for the Glu127 and Ile217 segment under specific molecular contexts.

**Figure 5.**
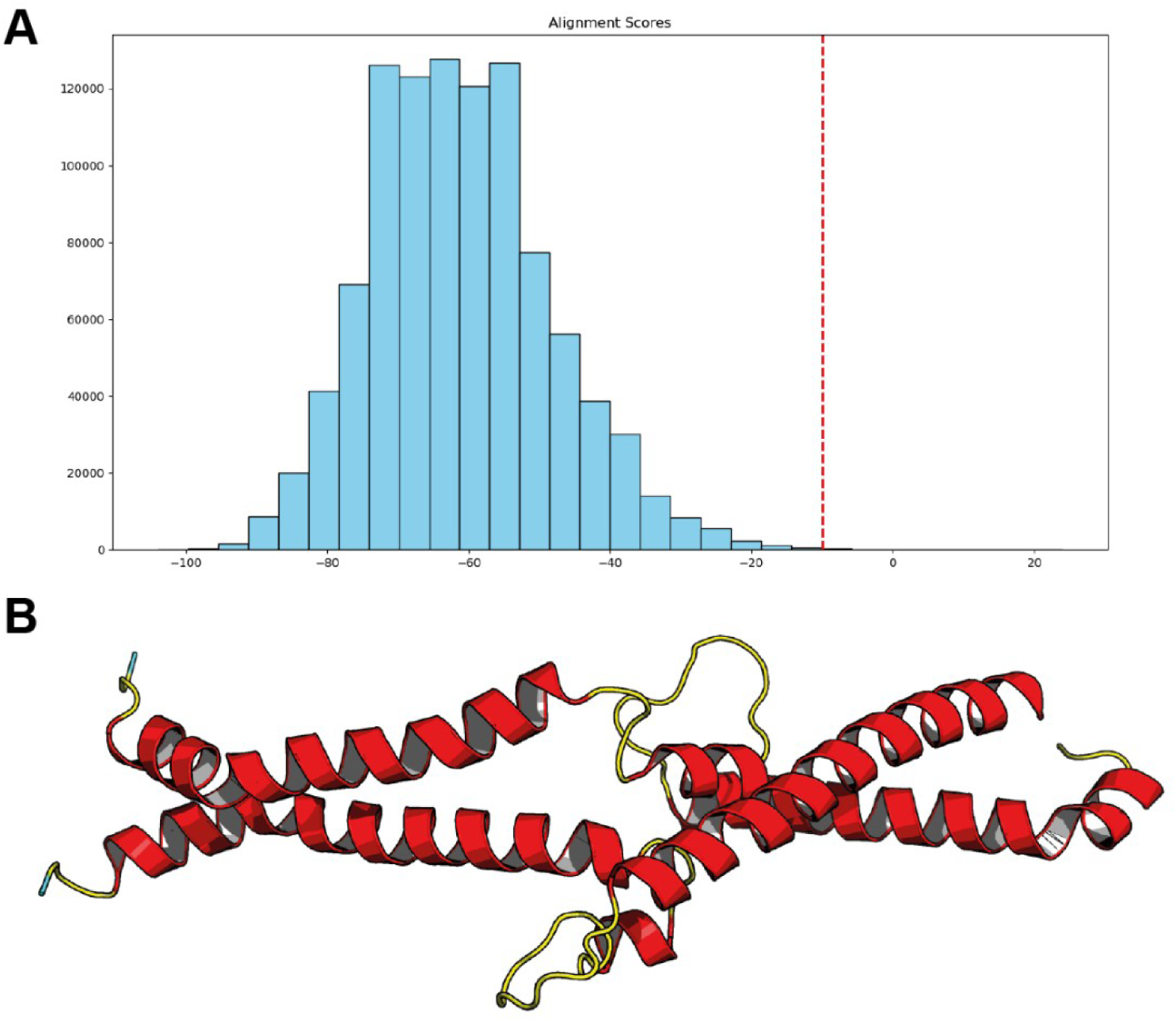
A segment of Rqc1p putatively adopts an HLH-like domain structure. **Panel A:** A shuffled sequence test for the significance of the alignment between the Glu127-Ile217 segment from Rqc1p and the MITF HLH domain (PDB: 7D8T). The histogram shows the scores for the best alignments between the two sequences when one of them is shuffled (repeated 10 ^3^), and the dotted red line corresponds to the actual alignment score between the two sequences. **Panel B:** A putative structure of the Rqc1p Glu127-Ile217 segment after being modelled using the MITF HLH structure as a template. The modelled structure was subjected to energy minimization and molecular dynamics consisting of 1 ns of NVT equilibration and 100 ns in NPT conditions.

### *S. cerevisiae* Rqc1p carries nuclear localization signals

Interestingly, many properties of Rqc1p uncovered during the course of this study are shared with specific protein families – in fact, conformational flexibility, intrinsically disordered regions, acid-rich segments, polybasic stretches potentially involved in electrostatic interactions, and protein dimerization domains are frequently observed in regulatory proteins associated with nuclear localization and nucleocytoplasmic functions (Homma et al., 2012; Frege & Uversky, 2015; Olsen et al., 2017; Tarczewska & Greb-Markiewicz, 2019). Based on these observations, we next investigated whether this protein presents nuclear localization signals (NLSs), which are short sequences of amino acid residues that can be act as tags to mediate the translocation of proteins found in the cytosol toward the nucleus through the nuclear pore complex (Lange et al., 2007).

Using DeepLoc 2.1 (Ødum et al., 2024), we verified that Rqc1p and its orthologs from human and mice, in fact, display cytosolic localization; also, while the latter two proteins also show a significant probability of nuclear localization – as initially expected –, Rqc1p did not, since its probability score (0.4957) remained below the prediction threshold (0.5014), though marginally (Table S5). However, this analysis further identified the presence of a putative NLS in Rqc1p, raising the possibility of this protein being translocated from the cytosol toward the nucleus under certain conditions. Notably, the residues contributing to the nuclear localization score of Rqc1p are clustered near to the N-terminal region of the protein – including its polybasic Lys/Arg stretch – which is highly conserved in eukaryotes (Figure 6A). Consistently, a second analysis using NucPred (Brameier et al., 2007) further supported this hypothesis, indicating a high nuclear localization score for Rqc1p (0.99), exceeding the recommended cutoff value (0.8), suggesting that Rqc1p is indeed likely to be translocated toward the nucleus (Table S5).

**Figure 6.**
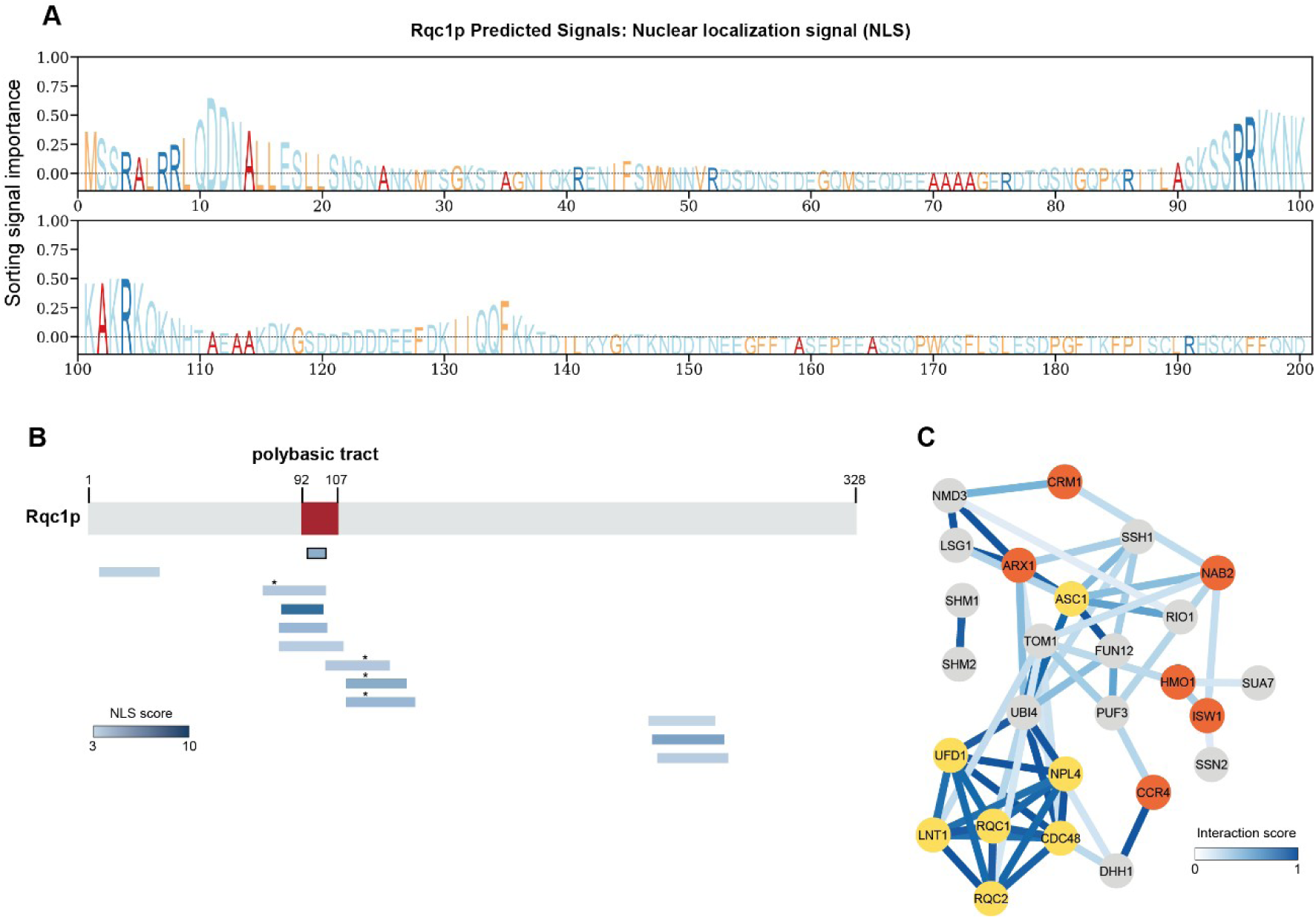
Rqc1p carries predicted nuclear localization features. **Panel A:** Prediction of subcellular localization scores and the presence of nuclear localization signals (NLS) along the Rqc1p N-terminal sequence based on DeepLoc 2.1. The logo-like plot represents residue-wise importance for localization prediction, where higher values indicate regions contributing to sorting signals. **Panel B:** Schematic representation of Rqc1p highlighting predicted NLS based on cNLS Mapper analysis. The polybasic tract is indicated and predicted monopartite (outlined box) and bipartite (boxes without a border) NLS sequences in the N-terminal are shown with their respective scores (blue gradient), where higher scores indicate stronger nuclear localization potential. Asterisks indicate serine residues annotated in the SGD database that may undergo phosphorylation as a post-translational modification. **Panel C:** Protein interaction network of Rqc1p based on physical interaction data retrieved from the SGD database and visualized in Cytoscape (Shannon et al., 2003), highlighting the associations with nuclear proteins involved in distinct aspects of nuclear metabolism (orange), in addition to components of the RQC pathway (yellow). Edge intensity represents interaction confidence.

Taken together, the results from DepLoc-2.1 and NucPred led us to conduct an additional, independent analysis to identify potential NLSs in Rqc1p using cNLS Mapper, which identifies and classifies NLSs canonically recognized by the importin α/β pathway (Kosugi et al., 2009). Surprisingly, this analysis determined multiple bipartite NLSs and one monopartite NLS in Rqc1p – notably, most of these NLSs are found within the polybasic tract of the protein (Figure 6B and Table S6), reinforcing the potential relevance of this region for the hypothetical translocation of Rqc1p to the nucleus.

Since the phosphorylation of amino acid residues within NLS motifs is a well-described regulatory mechanism that modulates protein nuclear import and subcellular localization (Harreman et al. 2004; Nardozzi et al., 2010), we next examined post-translational modification data available on SGD to determine whether any residues within the predicted Rqc1p NLSs are subject of this post-translational modification. Interestingly, the observation that there are two serine residues susceptible to phosphorylation – Ser64 and Ser80 – in four out of twelve bipartite NLSs indicated by cNLS Mapper (Figure 6B) strengthens the hypothesis that Rqc1p may undergo nucleocytoplasmic transport.

In fact, we found that several studies have reported Rqc1p physical association with nuclear proteins involved in diverse aspects of nuclear metabolism (Figure 6C) – these include direct interaction of Rqc1p with Arx1p, involved in pre-ribosome 60S association (Hung et al., 2008); Crm1p, involved in nuclear export (Kirli et al., 2015); Ccr4p, involved with mRNA poly(A) decay (Miller et al., 2018); Isw1p and Hmo1p, involved in chromatin remodeling (Babour et al., 2016; Marugesapillai et al. 2014); and Nab2p, involved in mRNA processing (Batisse et al., 2009), supporting the proposed nuclear localization of Rqc1p.

### *RQC1* transcription levels do not fluctuate under heat shock

To investigate whether the response of *S. cerevisiae* to heat stress involves alterations in gene expression levels of RQC complex proteins and proteins related to the HSR pathway, we decided to perform an analysis of the variation in transcript levels of genes of interest in yeast subjected to heat shock – i.e, an acute heat stress. To do so, we used publicly available data from studies that investigated the effect of this treatment on the transcriptome of *S. cerevisiae*. Principal component analysis (PCA) was then performed to determine the patterns of transcriptional variation between the experimental conditions of the studies and showed that, in all selected studies, there was distinct clustering among samples subjected to both experimental conditions – control or heat shock (Figure S4).

Then, we performed a differential gene expression analysis to evaluate the transcriptional response to heat shock across all conditions. Overall, the results revealed a clear time-dependent pattern of variation, with a higher number of differentially expressed genes (DEGs) detected under short-term heat shock exposure (Figure S5). To further investigate the influence of heat shock on the expression of genes in the RQC and HSR pathways, we selected a curated set of 28 genes, and found that most genes encoding proteins involved in the heat shock response are differentially expressed in all studies, especially under conditions of short-term heat shock (15, 20 and 30 min) (Figure 7A). Interestingly, with the exception of *ASC1* – always downregulated except under a heat shock condition conducted for 120 in one study – and *UFD1* – upregulated upon heat shock performed for 20 min –, all other genes in the RQC pathway exhibited unaltered transcriptional levels, suggesting that RQC proteins may be found in optimal cellular concentrations to allow a rapid response to the possible consequences of heat stress/heat shock on protein translation (Figure 7B).

**Figure 7.**
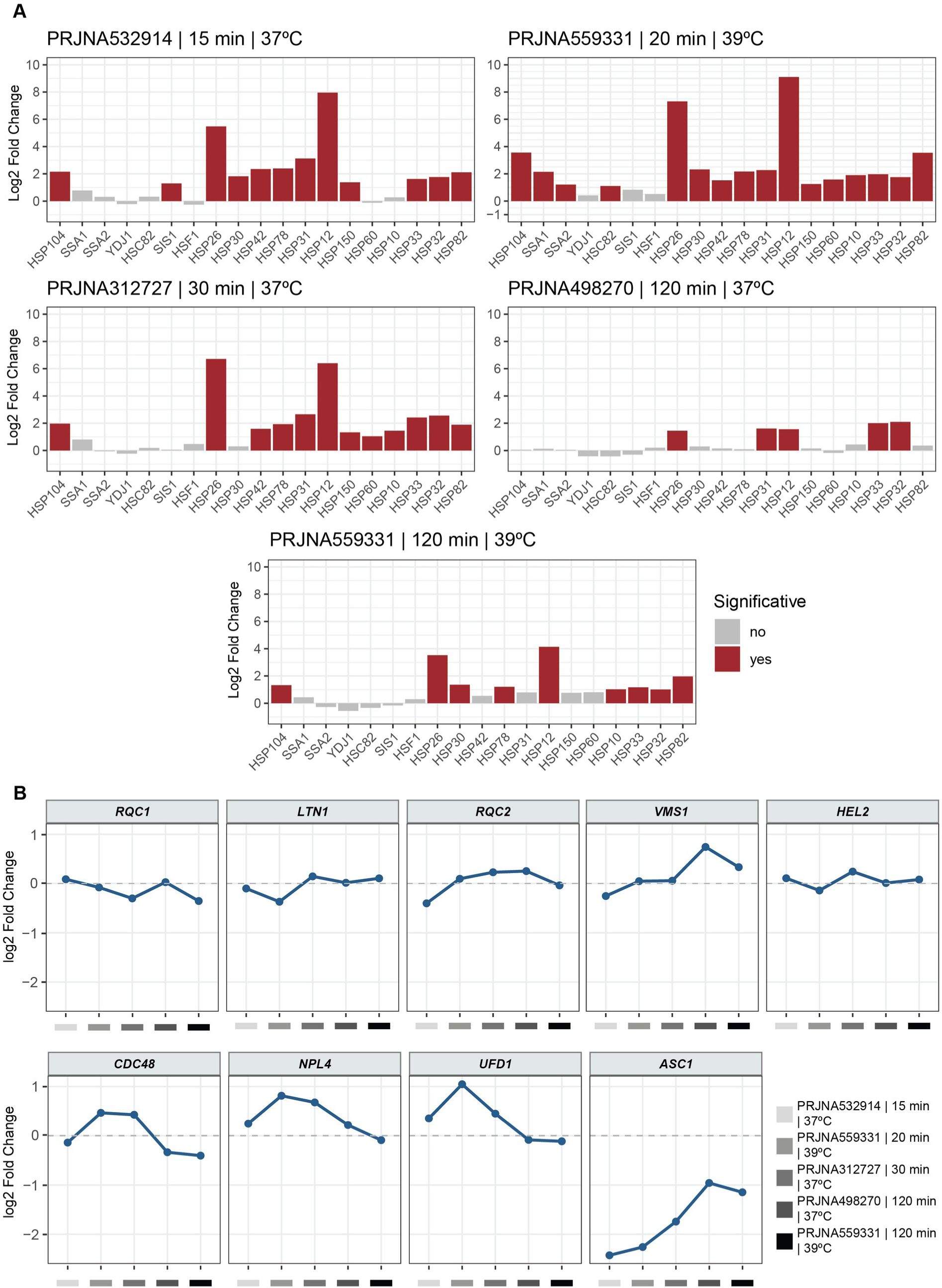
*RQC1* transcription levels do not fluctuate under heat shock. Temporal expression profiles of genes from the RQC and HSR pathways across RNA-seq datasets under heat shock conditions. **Panel A:** scatter plots showing expression changes across different times. **Panel B:** Barplot means genes are significant (red) and are not significant (grey). Genes were considered significantly differentially expressed when log2FoldChange ≥ |1| and adjusted *p*-value (padj) <0.05.

## Discussion

Proteome remodeling is characterized by the qualitative and quantitative reorganization of the set of proteins present within a cell in response to various biological scenarios such as cell differentiation, development, metabolic adaptation, environmental stress (whether thermal, oxidative, or osmotic), infection, and aging (Balch et al., 2008; Sui et al., 2020). This process is mediated by modulations in transcription, translation, and protein degradation, including protein synthesis quality control, selective oligo or polyubiquitination, and proteasomal or lysosomal degradation. In this manner, proteins unnecessary for the new biological conditions are disposed of, while those necessary for maintaining cellular fitness have their synthesis and/or stability increased (Dikic, 2017). One of the agents involved in this process is the proteostasis network, an integrated system that maintains a balance between protein synthesis, folding, trafficking, and degradation, ensuring the proteome’s functionality over time and promoting cellular fitness (Balch et al., 2008; Hipp et al., 2014). This network includes molecular chaperones, co-chaperones, ubiquitin-proteasome-dependent degradation systems – such as the RQC pathway –, autophagic pathways, translational surveillance mechanisms, and responses to proteotoxic stress, such as the unfolded protein response (UPR) and the heat shock response (HSR) (Anckar and Sistonen, 2011; Hartl et al., 2011; Walter & Ron, 2011; Brandman & Hedge, 2016).

The RQC pathway is functionally connected to HSR through the generation of proteotoxic stress when translational surveillance is persistently or dysfunctionally triggered (Defenouillère et al., 2013; Choe et al., 2016; Yonashiro et al., 2016). During polysomal collisions, the truncated nascent polypeptides are modified by the addition of Rqc2/NEMF-dependent CAT-tails. The accumulation of CAT-tailed polypeptides favors the formation of cytosolic protein aggregates, which sequester chaperones and compromise the basal capacity of the proteostasis network (Olzscha et al., 2011). This imbalance serves as an indirect signal for Hsf1p activation, the main HSR transcription factor, promoting the induction of transcription and translation of molecular chaperones and components of the degradation system. Interestingly, although *RQC1*-deficient cells exhibit Rqc2p-dependent CAT-tylation of newly-synthesized defective proteins (Yonashiro et al., 2016) – which would favor HSR activation and thus promote cytoprotection –, there is no observation of thermoadaptation in *rqc1*Δ cells as these mutants exhibit a marked sensitivity to heat stress in spot and growth kinetics assays (Figure 1). Although we cannot state whether or not HSF1 is activated in these mutants based on our available data, we demonstrated that *S. cerevisiae* thermotolerance strongly depends on the presence of Rqc1p, which was described as being part of the RQC pathway of this organism in earlier studies (Brandman et al., 2012). Rqc1p-deficient cells exhibit an accumulation of reporter proteins with CAT-tail, suggesting that this protein is involved in steps preceding or coinciding with the extraction of newly-synthesized defective proteins from the 60S subunit by the Cdc48p/Ufd1p/Npl4p complex (Defenouillère et al., 2016).

Interestingly, during the 2000s, homologs of Rqc1p were originally identified in mammals, and annotated as TCF25 or NULP1 as proteins from the bHLH family, and classified as possible regulators of embryonic tissue development (Olsson et al., 2002). Subsequently, it was demonstrated that human TCF25 can regulate the transcriptional activity of serum response factor (SRF) and is associated with the regulation of nuclear factor of activated T cells (NFAT) expression in cardiomyocytes during pathological cardiac hypertrophy (Cai et al., 2006; Zhang et al., 2020).

Multiple sequence alignment of *S. cerevisiae* Rqc1p and its homologs revealed the presence of two highly conserved regions in the yeast protein, being the first one located within the N-terminal domain, harboring the Lys/Arg polybasic tract, whereas and the second one is located further downstream and corresponds to the Tcf25 domain (Figure 2 A-B). Furthermore, once the identification of an HLH motif was not possible based on the available RQC complex-associated Rqc1p cryo-EM map by Li et al. (2025), we conducted structural predictions of unbound Rqc1p using molecular dynamics simulations, which suggested the presence of an HLH-like motif within its Tcf25 domain (Figure 5). In addition, intrinsically disordered regions were identified in its N- and C-terminal ends both by structure-based analysis and secondary structure prediction (Figure S2). Interestingly, many proteins harboring an HLH domain in fact display intrinsically disordered regions (Tarczewska and Greb-Markiewicz, 2019), which are not properly resolved by structural biology techniques relying on conformational homogeneity as they adopt multiple stable conformations in solution due to their large dependence on the cellular environment and its molecular interactions (Uversky, 2019).

Given that the findings obtained this study suggested that (i) Rqc1p may exhibit characteristics commonly associated with nuclear proteins, and that (ii) its mammalian orthologs have been described as transcription factors in human cells (Steen & Lindholm, 2008), we investigated the presence of localization signals and post-translational modification sites in the protein of *S. cerevisiae* through *in silico* investigation. Nucleocytoplasmic transport involves nuclear import and export activities, which are mediated by covalent modifications and interactions between nuclear localization signals (NLSs) and nuclear export signals (NESs) with specific proteins (Lange et al., 2007). Interestingly, analysis using NucPred indicated that *S. cerevisiae* Rqc1p may undergo nucleocytoplasmic relocalization (Table S5). In addition, assessment using DeepLoc 2.1 revealed an NLS associated with the N-terminal region of the sequence (Figure 6A). To validate these findings and enhance the robustness of the analyses, an additional *in silico* tool – cNLS Mapper – was employed to predict classical NLSs of the importin α/β pathway: this approach identified a monopartite NLS in Rqc1p, located at the N-terminal within the polybasic region, as well as several bipartite NLSs (Figure 6B and Table S6). Remarkably, initial studies characterizing mouse and human TCF25 identified a monopartite NLS (KKKKKQK) at position 127-133, which is similar to the NLS of the large T antigen of simian virus 40 (SV40) (Kalderon et al., 1984; Olsson et al., 2002; Cai et al., 2006).

Furthermore, the annotated phosphorylation sites for Rqc1p in SGD were mapped, revealing two serine residues within the predicted bipartite nuclear localization signals (NLSs) that are amenable to phosphorylation (Figure 6B and Table S6). Phosphorylation within or near an NLS is a well-established mechanism for regulating protein transport, as importin α-dependent phosphorylation around NLSs can mimic the inhibitory effect of acidic amino acids (Harreman et al., 2004), which leads to the hypothesis that, if Rqc1p translocation does occur, it would be a possible mechanism by which the process is conducted.

Finally, based on the finding that Rcp1p plays an important role in the HSR in *S. cerevisiae* (Figure 1), we investigated whether genes associated with the RQC complex pathway exhibit differential expression under this condition using publicly available RNA-seq data. The analysis of RNA-seq data from four independent studies revealed that genes associated with the classical HSR were differentially expressed, indicating that the response was captured in the experiments (Figure 7A). Most differentially expressed genes showed overexpression, with the greatest impact occurred between 15-20 minutes of heat shock treatment (Figure 7A). Prolonged treatment of 120 minutes led to a reduction in the expression of genes encoding HSPs, indicating that the stress response decreases over time (Figure 7A).

On the other hand, genes encoding proteins of the RQC pathway – including *RQC1* – did not show any difference in expression across conditions, except for *UFD1* and *ASC1* in some conditions (Figure 7B). One hypothesis to explain this observation is that the regulation of the core components of the RQC complex is independent of transcription, possibly via post-translational modification mechanisms. In fact, there is a proposed mechanism for the positive regulation of Rqc1p levels in the absence of Ltn1p, in which the former is presumably post-translationally degraded by the latter (Barros et al., 2021).

The absence of transcriptional changes in the wild-type strain does not preclude a role for *RQC1* in the HSR, particularly since *rqc1*Δ cells display a sensitivity phenotype that is rescued by gene reintroduction, as demonstrated in this study (Figure 2C). Nevertheless, the loss of cellular fitness resulting from the absence of *RQC1* appears to be a primary driver of the HSR in *S. cerevisiae*.

These findings open new avenues for future research on the potential role of Rqc1p as a nuclear-localized protein and the conditions that triggers its nuclear recruitment – in fact, subcellular localization assays in cells exposed to heat stress may clarify this hypothesis. Furthermore, biophysical assays, such as circular dichroism of individual domains, could verify their binding capacities to other proteins, and also nucleic acids, thereby elucidating the specific contributions of distinct Rqc1p regions. The search for mechanisms that explain the unexpected epistasis of *LTN1* over *RQC1* in heat stress situations (Figure 1B-D) may also explain whether ubiquitination in the absence of Rqc1p can prevent proper HSR activation upon heat stress, imbalancing thermoadaptation in *S. cerevisiae*.

In summary, the integrity of the RQC complex is essential for thermoadaptation in *S. cerevisiae*, as the absence of Rqc1p results in decreased thermotolerance. Structural analyses of Rqc1p indicate that this protein harbors a polybasic tract, an HLH-like structural motif, and intrinsically disordered regions, providing support for its longstanding classification within the bHLH family of eukaryotic regulatory proteins. In addition, the identification of putative NLSs suggests that Rqc1p may shuttle between the cytosol to the nucleus upon specific cellular contexts. Although Rqc1p is required for cytoprotection in supraoptimal temperature conditions, *RQC1* transcript levels remain unchanged under heat shock, suggesting that basal levels of Rqc1p are sufficient to fulfill its role in proteostasis during heat stress – this role may involve its well-established role in protein synthesis quality control, as well as a potential regulatory activity that has yet to be experimentally investigated.

## Acknowledgements

The authors thanks Ruth Maria Samaniego Valdez and Luis Eduardo Soares Netto for providing reagents, and Edson Alves Gomes for excellent technical support. ACPA was supported by a Master’s fellowship from Coordenação de Aperfeiçoamento de Pessoal de Nível Superior (CAPES), and FGCO by a Master’s fellowship from Fundação de Amparo à Pesquisa do Estado de Minas Gerais (FAPEMIG). LB was funded by CNPq (grant 308711/2022-0) and FAPEMIG (grant APQ-01898-22). EBT is grateful to Andréa Mara Macedo, Carlos Renato Machado and Mariana Torquato Quezado de Magalhães for academic support and access to instrumentation. Data processing and transcriptome analysis were performed on the Gloriosos computational clusters (Programa de Pós-Graduação em Bioinformática at Instituto de Ciências Biológicas, Universidade Federal de Minas Gerais).

## Conflict of interest

The authors declare that there are no conflicts of interest regarding the publication of this paper.

## Author contributions

ACPA: Conceived, designed and performed the experiments; analyzed the data; reviewed the manuscript.

FGCO: Designed and performed the experiments; analyzed the data; reviewed the manuscript. MMCL: Performed the experiments, analyzed the data; reviewed the manuscript.

AFC: Performed the experiments, analyzed the data; reviewed the manuscript. EMR: Performed the experiments, analyzed the data; reviewed the manuscript. GRF: Designed the experiments; analyzed the data; reviewed the manuscript.

MHB: Designed and performed the experiments; analyzed the data; reviewed the manuscript. LB: Designed and performed the experiments; analyzed the data; reviewed the manuscript.

EBT: Conceived, designed and performed the experiments; analyzed the data; drafted and reviewed the manuscript.

## Notes

### Competing Interest Statement

The authors have declared no competing interest.

## References

Abaeva, I. S., Bulakhov, A. G., Hellen, C. U. T., & Pestova, T. V. (2025). The ribosome-associated quality control factor TCF25 imposes K48 specificity on Listerin-mediated ubiquitination of nascent chains by binding and specifically orienting the acceptor ubiquitin. Genes & Development, 39, 617–633. 10.1101/gad.352389.124

Alagar Boopathy, L. R., Jacob-Tomas, S., Alecki, C., & Vera, M. (2022). Mechanisms tailoring the expression of heat shock proteins to proteostasis challenges. Journal of Biological Chemistry, 298(5), 101796. 10.1016/j.jbc.2022.101796

Alagar Boopathy, L. R., Beadle, E., Xiao, A. R., Garcia-Bueno Rico, A., Alecki, C., Garcia de-Andres, I., Edelmeier, K., Lazzari, L., Amiri, M., & Vera, M. (2023). The ribosome quality control factor Asc1 determines the fate of HSP70 mRNA on and off the ribosome. Nucleic Acids Research, 51(12), 6370–6388. 10.1093/nar/gkad338

Anckar, J., & Sistonen, L. (2011). Regulation of HSF1 function in the heat stress response: Implications in aging and disease. Annual Review of Biochemistry, 80(1), 1089–1115. 10.1146/annurev-biochem-060809-095203

Babour, A., Shen, Q., Dos-Santos, J., Murray, S., Gay, A., Challal, D., Fasken, M., Palancade, B., Corbett, A., Libri, D., Mellor, J., & Dargemont, C. (2016). The chromatin remodeler ISW1 is a quality control factor that surveys nuclear mRNP biogenesis. Cell, 167(5), 1201–1214.e15. 10.1016/j.cell.2016.10.048

Balch, W. E., Morimoto, R. I., Dillin, A., & Kelly, J. W. (2008). Adapting proteostasis for disease intervention. Science, 319(5865), 916–919. 10.1126/science.1141448

Balchin, D., Hayer-Hartl, M., & Hartl, F. U. (2016). In vivo aspects of protein folding and quality control. Science, 353(6294), aac4354. 10.1126/science.aac4354

Baler, R., Welch, W. J., & Voellmy, R. (1992). Heat shock gene regulation by nascent polypeptides and denatured proteins: hsp70 as a potential autoregulatory factor. Journal of Cell Biology, 117(6), 1151–1159. 10.1083/jcb.117.6.1151

Barros, G. C., Requião, R. D., Carneiro, R. L., Masuda, C. A., Moreira, M. H., Rossetto, S., Domitrovic, T., & Palhano, F. L. (2021). Rqc1 and other yeast proteins containing highly positively charged sequences are not targets of the RQC complex. Journal of Biological Chemistry, 296, 100586. 10.1016/j.jbc.2021.100586

Batisse, J., Batisse, C., Budd, A., Böttcher, B., & Hurt, E. (2009). Purification of nuclear poly(A)-binding protein Nab2 reveals association with the yeast transcriptome and a messenger ribonucleoprotein core structure. Journal of Biological Chemistry, 284(50), 34911–34917. 10.1074/jbc.m109.062034

Bengtson, M. H., & Joazeiro, C. A. P. (2010). Role of a ribosome-associated E3 ubiquitin ligase in protein quality control. Nature, 467, 470–473. 10.1038/nature09371

Best, R. B., Zhu, X., Shim, J., Lopes, P. E. M., Mittal, J., Feig, M., & MacKerell, A. D. (2012). Optimization of the Additive CHARMM All-Atom Protein Force Field Targeting Improved Sampling of the Backbone ϕ, ψ and Side-Chain χ_1_ and χ_2_ Dihedral Angles. Journal of Chemical Theory and Computation, *8*(9), 3257–3273. 10.1021/ct300400x

Bleicher, L., Lemke, N., & Garratt, R. C. (2011). Using amino acid correlation and community detection algorithms to identify functional determinants in protein families. PLoS ONE, 6(12), e27786. 10.1371/journal.pone.0027786

Bolivar, F., Rodriguez, R. L., Greene, P. J., Betlach, M. C., Heyneker, H. L., Boyer, H. W., Crosa, J. H., & Falkow, S. (1977). Construction and characterization of new cloning vehicles. II. A multipurpose cloning system. Gene, 2(2), 95–113. 10.1016/0378-1119(77)90000-2

Brachmann, C. B., Davies, A., Cost, G. J., Caputo, E., Li, J., Hieter, P., & Boeke, J. D. (1998). Designer deletion strains derived from Saccharomyces cerevisiae S288C: A useful set of strains and plasmids for PCR-mediated gene disruption and other applications. Yeast, 14(2), 115–132. 10.1002/(SICI)1097-0061(19980130)14:2

Brameier, M., Krings, A., & MacCallum, R. M. (2007). NucPred—Predicting nuclear localization of proteins. Bioinformatics, 23(9), 1159–1160. 10.1093/bioinformatics/btm066

Brandman, O., Stewart-Ornstein, J., Wong, D., Larson, A., Williams, Christopher C., Li, G.-W., Zhou, S., King, D., Shen, Peter S., Weibezahn, J., Dunn, Joshua G., Rouskin, S., Inada, T., Frost, A., & Weissman, Jonathan S. (2012). A ribosome-bound quality control complex triggers degradation of nascent peptides and signals translation stress. Cell, 151(5), 1042–1054. 10.1016/j.cell.2012.10.044

Brandman, O., & Hegde, R. S. (2016). Ribosome-associated protein quality control. Nature Structural & Molecular Biology, 23, 7–15. 10.1038/nsmb.3147

Braun, M. A., Costa, P. J., Crisucci, E. M., & Arndt, K. M. (2007). Identification of Rkr1, a Nuclear RING Domain Protein with Functional Connections to Chromatin Modification in *Saccharomyces cerevisiae*. Molecular and Cellular Biology, 27(8), 2800–2811. 10.1128/mcb.01947-06

Bukau, B., Weissman, J., & Horwich, A. (2006). Molecular chaperones and protein quality control. Cell, 125(3), 443–451. 10.1016/j.cell.2006.04.014

Cai, Z., Wang, Y., Yu, W., Xiao, J., Li, Y., Liu, L., Zhu, C., Tan, K., Deng, Y., Yuan, W., Liu, M., & Wu, X. (2006). hnulp1, a basic helix-loop-helix protein with a novel transcriptional repressive domain, inhibits transcriptional activity of serum response factor. Biochemical and Biophysical Research Communications, 343(3), 973–981. 10.1016/j.bbrc.2006.02.187

Cherry, J. M., Adler, C., Ball, C., Chervitz, S. A., Dwight, S. S., Hester, E. T., Jia, Y., Juvik, G., Roe, T., Schroeder, M., Weng, S., & Botstein, D. (1998). SGD: Saccharomyces genome database. Nucleic Acids Research, 26(1), 73–79. 10.1093/nar/26.1.73

Choe, Y. J., Park, S. H., Hassemer, T., Körner, R., Vincenz-Donnelly, L., Hayer-Hartl, M., & Hartl, F. U. (2016). Failure of RQC machinery causes protein aggregation and proteotoxic stress. Nature, 531, 191–195. 10.1038/nature16973

Creighton, T. E., (1993). Proteins: Structures and molecular properties (2^nd^ ed.). W. H. Freeman.

da Fonseca, N. J., Jr., Afonso, M. Q. L., de Oliveira, L. C., & Bleicher, L. (2019). A new method bridging graph theory and residue co-evolutionary networks for specificity determinant positions detection. Bioinformatics, 35(9), 1478–1485. 10.1093/bioinformatics/bty846

Danecek, P., Bonfield, J. K., Liddle, J., Marshall, J., Ohan, V., Pollard, M. O., Whitwham, A., Keane, T., McCarthy, S. A., Davies, R. M., & Li, H. (2021). Twelve years of SAMtools and BCFtools. GigaScience, 10(2), giab008. 10.1093/gigascience/giab008

Decker, C. J., & Parker, R. (2012). P-bodies and stress granules: Possible roles in the control of translation and mRNA degradation. Cold Spring Harbor Perspectives in Biology, 4, a012286. 10.1101/cshperspect.a012286

Defenouillère, Q., Yao, Y., Mouaikel, J., Namane, A., Galopier, A., Decourty, L., Doyen, A., Malabat, C., Saveanu, C., Jacquier, A., & Fromont-Racine, M. (2013). Cdc48-associated complex bound to 60S particles is required for the clearance of aberrant translation products. Proceedings of the National Academy of Sciences of the United States of America, 110(13), 5046–5051. 10.1073/pnas.1221724110

Defenouillère, Q., Zhang, E., Namane, A., Mouaikel, J., Jacquier, A., & Fromont-Racine, M. (2016). Rqc1 and Ltn1 prevent C-terminal alanine-threonine tail (CAT-tail)-induced protein aggregation by efficient recruitment of Cdc48 on stalled 60S subunits. Journal of Biological Chemistry, 291, 12245–12253. 10.1074/jbc.m116.722264

de Juan, D., Pazos, F., & Valencia, A. (2013). Emerging methods in protein co-evolution. Nature Reviews Genetics, 14, 249–261. 10.1038/nrg3414

Dikic, I. (2017). Proteasomal and autophagic degradation systems. Annual Review of Biochemistry, 86, 193–224. 10.1146/annurev-biochem-061516-044908

Dima, R. I., & Thirumalai, D. (2006). Determination of network of residues that regulate allostery in protein families using sequence analysis. Protein Science, 15(2), 258–268. 10.1110/ps.051767306

Dobin, A., Davis, C. A., Schlesinger, F., Drenkow, J., Zaleski, C., Jha, S., Batut, P., Chaisson, M., & Gingeras, T. R. (2013). STAR: Ultrafast universal RNA-seq aligner. Bioinformatics, 29(1), 15 –21. 10.1093/bioinformatics/bts635

Eddy, S.R. (2011). Accelerated profile HMM searches. PLoS Computational Biology, 7(10), e1002195. 10.1371/journal.pcbi.1002195

Edgar, R. C. (2004). MUSCLE: Multiple sequence alignment with high accuracy and high throughput. Nucleic Acids Research, 32(5), 1792–1797. 10.1093/nar/gkh340

Ewels, P., Magnusson, M., Lundin, S., & Käller, M. (2016). MultiQC: Summarize analysis results for multiple tools and samples in a single report. Bioinformatics, 32(19), 3047–3048. 10.1093/bioinformatics/btw354

Fonseca-Júnior, N. J., Afonso, M. Q. L., Oliveira, L. C., & Bleicher, L. (2018). PFstats: A network-based open tool for protein family analysis. Journal of Computational Biology, 25(5), 480–486. 10.1089/cmb.2017.0181

Frege, T., & Uversky, V. N. (2015). Intrinsically disordered proteins in the nucleus of human cells. Biochemistry and Biophysics Reports, 1, 33–51. 10.1016/j.bbrep.2015.03.003

Frischmeyer, P. A., van Hoof, A., O’Donnell, K., Guerrerio, A. L., Parker, R., & Dietz, H. C. (2002). An mRNA surveillance mechanism that eliminates transcripts lacking termination codons. Science, 295(5563), 2258–2261. 10.1126/science.1067338

Gasch, A.P., Spellman, P.T., Kao, C.M., Carmel-Harel, O., Eisen, M.B., Storz, G., Botstein, D., & Brown, P.O. (2000). Genomic expression programs in the response of yeast cells to environmental changes. Molecular Biology of the Cell, 11(12), 4241–4257. 10.1091/mbc.11.12.4241

Gietz, R. D., & Schiestl, R. H. (2007). High-efficiency yeast transformation using the LiAc/SS carrier DNA/PEG method. Nature Protocols, 2, 31–34. 10.1038/nprot.2007.13

Gomez-Pastor, R., Burchfiel, E. T., & Thiele, D. J. (2018). Regulation of heat shock transcription factors and their roles in physiology and disease. Nature Reviews Molecular Cell Biology, 19(1), 4–19. 10.1038/nrm.2017.73

Harreman, M. T., Kline, T. M., Milford, H. G., Harben, M. B., Hodel, A. E., & Corbett, A. H. (2004). Regulation of Nuclear Import by Phosphorylation Adjacent to Nuclear Localization Signals. Journal of Biological Chemistry, 279(20), 20613–20621. 10.1074/jbc.m401720200

Harju, S., Fedosyuk, H., & Peterson, K. R. (2004). Rapid isolation of yeast genomic DNA: Bust n’ Grab. BMC Biotechnology, 4, 8. 10.1186/1472-6750-4-8

Hanahan, D. (1983). Studies on transformation of Escherichia coli with plasmids. Journal of Molecular Biology, 166(4), 557–580. 10.1016/s0022-2836(83)80284-8

Hartl, F. U., Bracher, A., & Hayer-Hartl, M. (2011). Molecular chaperones in protein folding and proteostasis. Nature, 475(7356), 324–332. 10.1038/nature10317

Hipp, M. S., Park, S.-H., & Hartl, F. U. (2014). Proteostasis impairment in protein-misfolding and aggregation diseases. Trends in Cell Biology, 24(9), 506–514. 10.1016/j.tcb.2014.05.003

Høie, M. H., Kiehl, E. N., Petersen, B., Nielsen, M., Winther, O., Nielsen, H., Hallgren, J., & Marcatili, P. (2022). NetSurfP-3.0: Accurate and fast prediction of protein structural features by protein language models and deep learning. Nucleic Acids Research, 50(W1), W510–W515. 10.1093/nar/gkac439

Hodgkinson, C. A., Moore, K. J., Nakayama, A., Steingrímsson, E., Copeland, N. G., Jenkins, N. A., & Arnheiter, H. (1993). Mutations at the mouse microphthalmia locus are associated with defects in a gene encoding a novel basic-helix-loop-helix-zipper protein. Cell, 74(2), 395–404. 10.1016/0092-8674(93)90429-t

Homma, K., Fukuchi, S., Nishikawa, K., Sakamoto, S., & Sugawara, H. (2012). Intrinsically disordered regions have specific functions in mitochondrial and nuclear proteins. Molecular BioSystems, 8(1), 247–255. 10.1039/c1mb05208j

Hughes, A. L. (1994). The evolution of functionally novel proteins after gene duplication. Proceedings of the Royal Society B: Biological Sciences, 256(1346), 119–124. 10.1098/rspb.1994.0058

Hung, N.-J., Lo, K.-Y., Patel, S. S., Helmke, K., & Johnson, A. W. (2008). Arx1 is a nuclear export receptor for the 60S ribosomal subunit in yeast. Molecular Biology of the Cell, 19(2), 735–744. 10.1091/mbc.e07-09-0968

Inada, T. (2013). Quality control systems for aberrant mRNAs induced by aberrant translation elongation and termination. Biochimica et Biophysica Acta (BBA) – Gene Regulatory Mechanisms, 1829(6–7), 634–642. 10.1016/j.bbagrm.2013.02.004

Isken, O., & Maquat, L. E. (2007). Quality control of eukaryotic mRNA: Safeguarding cells from abnormal mRNA function. Genes & Development, 21, 1833–1856. 10.1101/gad.1566807

Jo, S., Kim, T., Iyer, V. G., & Im, W. (2008). CHARMM-GUI: A web-based graphical user interface for CHARMM. Journal of Computational Chemistry, 29(11), 1859–1865. 10.1002/jcc.20945

Joazeiro, C. A. P. (2019). Mechanisms and functions of ribosome-associated protein quality control. Nature Reviews Molecular Cell Biology, 20, 368–383. 10.1038/s41580-019-0118-2

Jones, S. (2004). An overview of the basic helix-loop-helix proteins. Genome Biology, 5, 226. 10.1186/gb-2004-5-6-226

Kalderon, D., Roberts, B. L., Richardson, W. D., & Smith, A. E. (1984). A short amino acid sequence able to specify nuclear location. Cell, 39(3), 499–509. 10.1016/0092-8674(84)90457-4

Kırlı, K., Karaca, S., Dehne, H. J., Samwer, M., Pan, K. T., Lenz, C., Urlaub, H., & Görlich, D. (2015). A deep proteomics perspective on CRM1-mediated nuclear export and nucleocytoplasmic partitioning. eLife, 4, e11466. 10.7554/eLife.11466

Kosugi, S., Hasebe, M., Tomita, M., & Yanagawa, H. (2009). Systematic identification of cell cycle-dependent yeast nucleocytoplasmic shuttling proteins by prediction of composite motifs. Proceedings of the National Academy of Sciences of the United States of America, 106 (25), 10171–10176. 10.1073/pnas.0900604106

Kuroha, K., Zinoviev, A., Hellen, C. U. T., & Pestova, T. V. (2018). Release of ubiquitinated and non-ubiquitinated nascent chains from stalled mammalian ribosomal complexes by ANKZF1 and Ptrh1. Molecular Cell, 72(2), 286–302.e8. 10.1016/j.molcel.2018.08.022

Kuznetsov, D., Tegenfeldt, F., Manni, M., Seppey, M., Berkeley, M., Kriventseva, E. V., & Zdobnov, E. M. (2023). OrthoDB v11: Annotation of orthologs in the widest sampling of organismal diversity. Nucleic Acids Research, 51(D1), D445–D451. 10.1093/nar/gkac998

Kwolek-Mirek, M., & Zadrag-Tecza, R. (2014). Comparison of methods used for assessing the viability and vitality of yeast cells. FEMS Yeast Research, 14(7), 1068–1079. 10.1111/1567-1364.12202

Lange, A., Mills, R. E., Lange, C. J., Stewart, M., Devine, S. E., & Corbett, A. H. (2007). Classical nuclear localization signals: Definition, function, and interaction with importin α. Journal of Biological Chemistry, 282(8), 5101–5105. 10.1074/jbc.r600026200

Li, W., Scheel, T., & Shen, P. S. (2025). Mechanism of nascent chain removal by the ribosome-associated quality control complex. Nature Communications, 16, 5792. 10.1038/s41467-025-61235-w

Liao, Y., Smyth, G. K., & Shi, W. (2014). featureCounts: An efficient general purpose program for assigning sequence reads to genomic features. Bioinformatics, 30(7), 923–930. 10.1093/bioinformatics/btt656

Love, M. I., Huber, W., & Anders, S. (2014). Moderated estimation of fold change and dispersion for RNA-seq data with DESeq2. Genome Biology, 15(12), 550. 10.1186/s13059-014-0550-8

Luscombe, N. M., Austin, S. E., Berman, H. M., & Thornton, J. M. (2000). An overview of the structures of protein–DNA complexes. Genome Biology, 1(1), Reviews001. 10.1186/gb-2000-1-1-reviews001

Lyumkis, D., Oliveira dos Passos, D., Tahara, E. B., Webb, K., Bennett, E. J., Vinterbo, S., Potter, C. S., Carragher, B., & Joazeiro, C. A. P. (2014). Structural basis for translational surveillance by the large ribosomal subunit-associated protein quality control complex. Proceedings of the National Academy of Sciences of the United States of America, 111(45), 15981–15986. 10.1073/pnas.1413882111

Madeira, F., Madhusoodanan, N., Lee, J., Eusebi, A., Niewielska, A., Tivey, A. R. N., Lopez, R., & Butcher, S. (2024). The EMBL-EBI Job Dispatcher sequence analysis tools framework in 2024. Nucleic Acids Research, 52(W1), W521–W525. 10.1093/nar/gkae241

Marks, D. S., Hopf, T. A., & Sander, C. (2012). Protein structure prediction from sequence variation. Nature Biotechnology, 30, 1072–1080. 10.1038/nbt.2419

Masser, A. E., Kang, W., Roy, J.; Kaimal, J. M., Quintana-Cordero, J., Friedländer, M. R., & Andréasson, C. (2019). Cytoplasmic protein misfolding titrates Hsp70 to activate nuclear Hsf1. eLife, 8, e47791. 10.7554/eLife.47791

Matsuo, Y., Ikeuchi, K., Saeki, Y., Iwasaki, S., Schmidt, C., Udagawa, T., Sato, F., Tsuchiya, H., Becker, T., Tanaka, K., Ingolia, N. T., Beckmann, R., & Inada, T. (2017). Ubiquitination of stalled ribosome triggers ribosome-associated quality control. Nature Communications, 8, 159. 10.1038/s41467-017-00188-1

Miller, J. E., Zhang, L., Jiang, H., Li, Y., Pugh, B. F., & Reese, J. C. (2018). Genome-wide mapping of decay factor–mRNA interactions in yeast identifies nutrient-responsive transcripts as targets of the deadenylase Ccr4. G3: Genes, Genomes, Genetics, *8*(1), 315–330. 10.1534/g3.117.300415

Monaghan, L., Longman, D., & Cáceres, J. F. (2023). Translation-coupled mRNA quality control mechanisms. The EMBO Journal, 42, EMBJ2023114378. 10.15252/embj.2023114378

Monod, J. (1949). The growth of bacterial cultures. Annual Review of Microbiology, 3, 371–394. 10.1146/annurev.mi.03.100149.002103

Morimoto, R. I. (1998). Regulation of the heat shock transcriptional response: cross talk between a family of heat shock factors, molecular chaperones, and negative regulators. Genes & Development, 12(24), 3788–3796. 10.1101/gad.12.24.3788

Morimoto, R. I. (2008). Proteotoxic stress and inducible chaperone networks in neurodegenerative disease and aging. Genes & Development, 22(11), 1427–1438. 10.1101/gad.1657108

Murre, C., Bain, G., van Dijk, M. A., Engel, I., Furnari, B. A., Massari, M. E., Matthews, J. R., Quong, M. W., Rivera, R. R., & Stuiver, M. H. (1994). Structure and function of helix-loop-helix proteins. Biochimica et Biophysica Acta (BBA) - Gene Structure and Expression, 1218(2), 129 –135. 10.1016/0167-4781(94)90001-9

Murre, C. (2019). Helix–loop–helix proteins and the advent of cellular diversity: 30 years of discovery. Genes & Development, 33, 6–25. 10.1101/gad.320663.118

Murugesapillai, D., McCauley, M. J., Huo, R., Nelson Holte, M. H., Stepanyants, A., Maher, L. J., Israeloff, N. E., & Williams, M. C. (2014). DNA bridging and looping by HMO1 provides a mechanism for stabilizing nucleosome-free chromatin. Nucleic Acids Research, 42(14), 8996–9004. 10.1093/nar/gku635

Nardozzi, J. D., Lott, K., & Cingolani, G. (2010). Phosphorylation meets nuclear import: A review. Cell Communication and Signaling, 8(1), 32. 10.1186/1478-811X-8-32

Newman, M. E. J., & Girvan, M. (2004). Finding and evaluating community structure in networks. Physical Review E, 69, 026113. 10.1103/physreve.69.026113

Nuño-Cabanes, C., Ugidos, M., Tarazona, S., Martín-Expósito, M., Ferrer, A., Rodríguez-Navarro, S., & Conesa, A. (2020). A multi-omics dataset of heat shock response in the yeast RNA binding protein Mip6. Scientific Data, 7, 69. 10.1038/s41597-020-0412-z

Ødum, M. T., Teufel, F., Thumuluri, V., Almagro Armenteros, J. J., Johansen, A. R., Winther, O., & Nielsen, H. (2024). DeepLoc 2.1: Multi-label membrane protein type prediction using protein language models. Nucleic Acids Research, 52(W1), W215–W220. 10.1093/nar/gkae237

Olsen, J. G., Teilum, K., & Kragelund, B. B. (2017). Behaviour of intrinsically disordered proteins in protein–protein complexes with an emphasis on fuzziness. Cellular and Molecular Life Sciences, 74, 3175–3183. 10.1007/s00018-017-2560-7

Olsson, M., Durbeej, M., Ekblom, P., & Hjalt, T. (2002). Nulp1, a novel basic helix-loop-helix protein expressed broadly during early embryonic organogenesis and prominently in developing dorsal root ganglia. Cell and Tissue Research, 308, 361–370. 10.1007/s00441-002-0544-9

Olzscha, H., Schermann, S. M., Woerner, A. C., Pinkert, S., Hecht, M. H., Tartaglia, G. G., Vendruscolo, M., Hayer-Hartl, M., Hartl, F. U., & Vabulas, R. M. (2011). Amyloid-like aggregates sequester numerous metastable proteins with essential cellular functions. Cell, 144(1), 67–78. 10.1016/j.cell.2010.11.050

Pabo, C. O., & Sauer, R. T. (1992). Transcription factors: Structural families and principles of DNA recognition. Annual Review of Biochemistry, 61, 1053–1095. 10.1146/annurev.bi.61.070192.005201

Peffer, S., Gonçalves, D., & Morano, K. A. (2019). Regulation of the Hsf1-dependent transcriptome via conserved bipartite contacts with Hsp70 promotes survival in yeast. Journal of Biological Chemistry, 294(32), 12191–12202. 10.1074/jbc.ra119.008822

Phillips, J. C., Hardy, D. J., Maia, J. D. C., Stone, J. E., Ribeiro, J. V., Bernardi, R. C., Buch, R., Fiorin, G., Hénin, J., Jiang, W., McGreevy, R., Melo, M. C. R., Radak, B. K., Skeel, R. D., Singharoy, A., Wang, Y., Roux, B., Aksimentiev, A., Luthey-Schulten, Z., & Kalé, L. V. (2020). Scalable molecular dynamics on CPU and GPU architectures with NAMD. The Journal of Chemical Physics, 153(4), 044130. 10.1063/5.0014475

Poveda-Huertes, D., Matic, S., Marada, A., Habernig L., Licheva, M., Myketin, L., Gilsbach, R., Tosal-Castano, S., Papinski, D., Mulica, P., Kretz, O., Küçükköse, C., Taskin, A. A., Hein, L., Kraft, C.; Büttner, S., Meisinger, C., & Vögtle, F.-N. (2020). An early mtUPR: Redistribution of the nuclear transcription factor Rox1 to mitochondria protects against intramitochondrial proteotoxic aggregates. Molecular Cell, 77(1), 180–188. 10.1016/j.molcel.2019.09.026

Richter, K., Haslbeck, M., & Buchner, J. (2010). The heat shock response: Life on the verge of death. Molecular Cell, 40(2), 253–266. 10.1016/j.molcel.2010.10.006

Robinson, K. A., & Lopes, J. M. (2000). Saccharomyces cerevisiae basic helix–loop–helix proteins regulate diverse biological processes. Nucleic Acids Research, 28(7), 1499–1505. 10.1093/nar/28.7.1499

Šali, A., & Blundell, T. L. (1993). Comparative protein modelling by satisfaction of spatial restraints. Journal of Molecular Biology, 234(3), 779–815. 10.1006/jmbi.1993.1626

Sampaio-Marques, B., & Ludovico, P. (2018). Linking cellular proteostasis to yeast longevity. FEMS Yeast Research, 18(5), foy043. 10.1093/femsyr/foy043

Santiago, A. M., Gonçalves, D. L., & Morano, K. A. (2020). Mechanisms of sensing and response to proteotoxic stress. Experimental Cell Research, 395(2), 112240. 10.1016/j.yexcr.2020.112240

Schrödinger, LLC. (2026). The PyMOL Molecular Graphics System (Version 3.0).

Shannon, P. et al. (2003). Cytoscape: a software environment for integrated models of biomolecular interaction networks. Genome Research, 13(11), 2498–2504. 10.1101/gr.1239303

Sitron, C. S., Park, J. H., & Brandman, O. (2017). Asc1, Hel2, and Slh1 couple translation arrest to nascent chain degradation. RNA, *23*(5), 798–810. 10.1261/rna.060897.117

Staudinger, J., Perry, M., Elledge, S. J., & Olson, E. N. (1993). Interactions among vertebrate helix-loop-helix proteins in yeast using the two-hybrid system. Journal of Biological Chemistry, 268(7), 4608–4611. 10.1016/s0021-9258(18)53440-2

Steen, H., & Lindholm, D. (2008). Nuclear localized protein-1 (Nulp1) increases cell death of human osteosarcoma cells and binds the X-linked inhibitor of apoptosis protein. Biochemical and Biophysical Research Communications, 366(2), 432–437. 10.1016/j.bbrc.2007.11.146

Sui, X., Pires, D. E. V., Ormsby, A. R., Cox, D., Nie, S., Vecchi, G., Vendruscolo, M., Ascher, D. B., Reid, G. E., & Hatters, D. M. (2020). Widespread remodeling of proteome solubility in response to different protein homeostasis stresses. Proceedings of the National Academy of Sciences of the United States of America, 117(5), 2422–2431. 10.1073/pnas.1912897117

Tarczewska, A., & Greb-Markiewicz, B. (2019). The significance of the intrinsically disordered regions for the functions of the bHLH transcription factors. International Journal of Molecular Sciences, 20(21), 5306. 10.3390/ijms20215306

Tsuchiya, H., Ohtake, F., Arai, N., Kaiho, A., Yasuda, S., Tanaka, K., & Saeki, Y. (2017). In vivo ubiquitin linkage-type analysis reveals that the Cdc48-Rad23/Dsk2 axis contributes to K48-linked chain specificity of the proteasome. Molecular Cell, 66(4), 488–502.e7. 10.1016/j.molcel.2017.04.024

UniProt Consortium. (2025). UniProt: The Universal Protein Knowledgebase in 2025. Nucleic Acids Research, 53(D1), D609–D617. 10.1093/nar/gkae1010

Uversky, V. N. (2019). Intrinsically disordered proteins and their “mysterious” (meta)physics. Frontiers in Physics, 7. 10.3389/fphy.2019.00010

Valdez, R. M. S. (2018). *Impact of translational disruption on the metabolic fate of newly synthesized polypeptides in Saccharomyces cerevisiae* (Master’s thesis, Universidade Federal de Minas Gerais).

van Hoof, A., Frischmeyer, P. A., Dietz, H. C., & Parker, R. (2002). Exosome-mediated recognition and degradation of mRNAs lacking a termination codon. Science, 295(5563), 2262–2264. 10.1126/science.1067272

Vihervaara, A., Duarte, F.M., Lis, J.T. (2018) Molecular mechanisms driving transcriptional stress responses. Nature Reviews Genetics, 19(6), 385–397. 10.1038/s41576-018-0001-6

Walsh, J. B. (2003). Population-genetic models of the fates of duplicate genes. Genetica, 118, 279–294. 10.1023/A:1024194802441

Walter, P., & Ron, D. (2011). The unfolded protein response: From stress pathway to homeostatic regulation. Science, 334(6059), 1081–1086. 10.1126/science.1209038

Waterhouse, A. M., Procter, J. B., Martin, D. M. A., Clamp, M., & Barton, G. J. (2009). Jalview version 2: A multiple sequence alignment editor and analysis workbench. Bioinformatics, 25(9), 1189–1191. 10.1093/bioinformatics/btp033

Winz, M. L., Peil, L., Turowski, T. W., Rappsilber, J., & Tollervey, D. (2019). Molecular interactions between Hel2 and RNA supporting ribosome-associated quality control. Nature Communications, 10, 563. 10.1038/s41467-019-08382-z

Yonashiro, Ryo, Tahara, E. B., Bengtson, M. H., Khokhrina, M., Lorenz, H., Chen, K.-C., Kigoshi-Tansho, Y., Savas, J. N., Yates, J. R., III, Kay, S. A., Craig, E. A., Mogk, A., Bukau, B., & Joazeiro, C. A. P. (2016). The Rqc2/Tae2 subunit of the ribosome-associated quality control (RQC) complex marks ribosome-stalled nascent polypeptide chains for aggregation. eLife, 5, e11794. 10.7554/elife.11794

Zhang, X., Lei, F., Wang, X., Deng, K., Ji, Y., Zhang, Y., Li, H., Zhang, X., Lu, Z., & Zhang, P. (2020). NULP1 Alleviates Cardiac Hypertrophy by Suppressing NFAT3 Transcriptional Activity. Journal of the American Heart Association, 9(16), e016419. 10.1161/jaha.120.016419

Zeng, Q., Wan, Y., Zhu, P., Zhao, M., Jiang, F., Chen, J., Tang, M., Zhu, X., Li, Y., Zha, H., Wang, Y., Hu, M., Mo, X., Zhang, Y., Chen, Y., Chen, Y., Ye, X., Bodmer, R., Ocorr, K., Jiang, Z., Zhuang, J., Yuan, W., & Wu, X. (2018). The bHLH protein Nulp1 is essential for femur development via acting as a cofactor in Wnt signaling in Drosophila. Current Molecular Medicine, 17(7), 509–517. 10.2174/1566524018666180212145714

Zheng, X., Krakowiak, J., Patel, N., Beyzavi, A., & Pincus, D. (2016). Dynamic control of Hsf1 during heat shock by a chaperone switch and phosphorylation. eLife, 5:e18638. 10.7554/elife.18638

