## Supplementary Information for "Structural and functional insights into yeast Rqc1p, a protein required for thermotolerance with potential nuclear localization"

**Running title: Insights into *S. cerevisiae* Rqc1p**

Amanda Cristina Pereira-Antônio[1]; Frederico Gabriel de Carvalho Oliveira[1]; Marcelo Mesa Costa-Lima[2]; Aysllan Fernandes Coelho[1]; Erik Marques Rodrigues[1]; Glória Regina Franco[1]; Mario Henrique de Barros[2]; Lucas Bleicher[1]; Erich Birelli Tahara[1,*]

[1]Departamento de Bioquímica e Imunologia, Instituto de Ciências Biológicas, Universidade Federal de Minas Gerais – Belo Horizonte, MG, Brazil

[2]Departamento de Microbiologia, Instituto de Ciências Biomédicas, Universidade de São Paulo – São Paulo, SP, Brazil

**Table S1: Yeast strains used in this study.**

| Strain | Genotype | Reference |
|---|---|---|
| BY4741 | *MATa his3Δ1 leu2Δ0 met15Δ0 ura3Δ0* | BRACHMANN et al., 1998 |
| *ltn1*Δ | *MATa: his3Δ1; leu2Δ0; met15Δ0; ura3Δ0; ltn1::KanMX6* | BRAUcN et al., 2007 |
| *rqc1*Δ | *MATa: his3Δ1; leu2Δ0; met15Δ0; ura3Δ0; rqc1::KanMX6* | BRANDMAN et al., 2012 |
| *rqc1*Δ*ltn1*Δ | *MATa: his3Δ1; leu2Δ0; met15Δ0; ura3Δ0 rqc1::KanMX6; ltn1Δ::His3MX6* | VALDEZ, 2018 |
| *rqc1*ΔRqc1p | *MATa: his3Δ1; leu2Δ0; met15Δ0; ura3Δ0 rqc1::KanMX6* YCplac111-Rqc1p$^{1-723}$ | This study |
| *rqc1*ΔRqc1p$^{1-327}$ | *MATa: his3Δ1; leu2Δ0; met15Δ0; ura3Δ0 rqc1::KanMX6* YCplac111-*rqc1*ΔRqc1p$^{1-327}$ | This study |
| *rqc1*ΔRqc1p$^{328-723}$ | *MATa: his3Δ1; leu2Δ0; met15Δ0; ura3Δ0rqc1::KanMX6* YCplac111-Rqc1p$^{328-723}$ | This study |

**Table S2. General features of orthologous protein sequences from selected model organisms retrieved from Uniprot.**

| Accession | Organism | Status review | Annotation score | Protein existence | Amino acids |
|---|---|---|---|---|---|
| Q05468 | *Saccharomyces cerevisiae* | Swiss-Prot | 4/5 | Evidence at protein level | 723 |
| Q9BQ70 | *Homo sapiens* | Swiss-Prot | 5/5 | Evidence at protein level | 676 |
| Q8R3L2 | *Mus musculus* | Swiss-Prot | 5/5 | Evidence at protein level | 676 |
| A0A8V0Y0J9 | *Gallus gallus* | TrEMBL | 1/5 | Evidence at protein level | 661 |
| Q68EJ4 | *Danio rerio* | TrEMBL | 1/5 | Evidence at protein level | 650 |
| A3KMT4 | *Xenopus laevis* | TrEMBL | 1/5 | Evidence at transcript level | 675 |
| Q9Y109 | *Drosophila melanogaster* | TrEMBL | 2/5 | Evidence at transcript level | 702 |
| P90919 | *Caenorhabiditis elegans* | TrEMBL | 1/5 | Evidence at protein level | 647 |
| Q8T2A4 | *Dictyostelium discoideum* | Swiss-Prot | 3/5 | Inferred from homology | 740 |

| | | | | | |
|---|---|---|---|---|---|
| O80734 | *Arabidopsis thaliana* | TrEMBL | 1/5 | Evidence at protein level | 627 |

**Table S3: Primers used for cloning and sequencing.** Restriction sites recognized by the restriction enzymes are shown in bold.

| Primer | Sequence (5’ -> 3’) | Application |
|---|---|---|
| pRQC1_FW_(HindIII) | ATGC**AAGCTT**CTTTTGAACTTCAAGGACTTCGCTGC | Promoter cloning |
| pRQC1_RV_(PstI) | GTAC**CTGCAG**GTTCGAGTACTTTACAAATATATTTAGATGATTCAACGACC | Promotor cloning |
| RQC1_FW_ALL_(PstI) | ATCG**CTGCAG**ATGAGCTCTAGAGCATTAAGGAGATTAC | Full-length and N-terminal cloning |
| RQC1_RV_ALL_(KpnI) | GTAC**GGTACC**TTATTTATCGTCATCGTCTTTATAATCACCCTCATTTTCATTTGACTCTTCATTTTCATG | Full-length and C-terminal cloning |
| RQC1_NT_RV_(KpnI) | TACG**GGTACC**TTATTTATCGTCATCGTCTTTATAATCATAAAAAGAAATGAATTTTTGCCACTGACTG | N-terminal cloning |
| RQC1_CT_FW_(PstI) | ATCG**CTGCAG**ATGAAGTTCGAACCTTTAAATTCCGACCTGAGC | C-terminal cloning |
| tRQC1_FW_(KpnI) | ATCG**GGTACC**TAAGAAATGTTGCAGCTCATTTTCTAGTACGC | Terminator cloning |
| tRQC1_RV_(EcoRI) | CGAT**GAATTC**CGGACTTGGAAGGACAAACAATGC | Terminator cloning |
| Seq_1 | CTCGTAAGTTGTCGCGAAACGC | Sequencing of YCplac111 + Rqc1p constructs |
| Seq_2 | CGAGGCTGCAAAAGACAAGGG | Sequencing of full-length and N-terminal constructs |
| Seq_3 | CCAAAACGATGTCAGTCAGTGGC | Sequencing of full-length construct |
| Seq_4 | GAATAAATGAGATGAGTTCCGCCCG | Sequencing of full-length and C-terminal constructs |
| Seq_5 | CGCCAGCTGGCGTAATAGCGAAG | Sequencing of full-length and N-terminal constructs |

**Table S4: Description of sequencing data obtained from the SRA.**

| **Bioproject** | **Sequencing platform** | **Library depth** | **Read length** | ***S. cerevisiae* strain** | **Growth temperature** | **Duration of heat shock treatment** | **Reference** |
|---|---|---|---|---|---|---|---|
| PRJNA532914 | Illumina HiSeq 4000 | 22,8 - 31,5M | 150 bp | WT BY4741 | 30ºC (control) and 37ºC (treated) | 15 min | Peffer, 2019 |
| PRJNA559331 | Illumina HiSeq 2000 | 26,8 - 33,3M | 101 bp | WT BY4741 | 30ºC (control) and 39ºC (treated) | 20 and 120 min | Nuño-Cabanes, 2020 |
| PRJNA312727 | Illumina HiSeq 2500 | 8,3 - 10,6M | 101 bp | WT CAY1015 | 25ºC (control) and 37ºC (treated) | 30 min | Masser, 2019 |
| PRJNA498270 | Illumina HiSeq 2500 | 9,9 - 12,0M | 50 bp | WT YPH499 | 23ºC (control) and 37ºC (treated) | 120 min | Poveda-Huertes, 2020 |

**Table S5: Predicted subcellular localization of Rqc1p and its homologous proteins using DeepLoc-2.1 and NucPred.** Localization prediction scores for TCF25 family proteins are shown. For DeepLoc-2.1, probabilities >0.4761 indicate likely cytoplasmic localization and >0.5014 indicate likely nuclear localization. Green cells highlight predictions, with color intensity indicating the certainty of the prediction. For NucPred, scores ≥ 0.8 indicate proteins predicted to spend time in the nucleus.

| | **DeepLoc-2.1** | | **NucPred** |
|---|---|---|---|
| Specie | Cytoplasm | Nucleus | Nucleus |
| *S. cerevisiae* | 0.6625 | 0.4957 | 0.99 |
| *H. sapiens* | 0.6098 | 0.5215 | 0.92 |
| *M. musculus* | 0.6081 | 0.5295 | 0.90 |

**Table S6: Predicted nuclear localization signals (NLSs) identified in Rqc1p using cNLS Mapper.** Putative monopartite and bipartite NLS were identified by prediction using cNLS Mapper with a cutoff score of 3.0 and screening across the full protein sequence. The table lists the NLS class, amino acid positions and prediction score for each identified sequence. Higher scores indicate a greater probability of nuclear localization, with scores of 8-10, 7-8, and 3-5 correspond to predominantly nuclear, partially nuclear, and both nuclear/cytoplasmatic localization, respectively. Underlined residue indicates phosphorylation sites annotated in the SGD database.

| NLS class | Position | Sequence | Score |
|---|---|---|---|
| Monopartite | 94 - 102 | SRRKKNKKA | 5 |
| Bipartite | 5 - 31 | ALRRLQDDNALLESLSSNSANMKTSGK | 3.3 |
| | 75 - 102 | ERDTQSNGQPKRITLASKSSRRKNKKAK | 3.7 |
| | 83 - 101 | QPKRITLASKSSRRKNKKK | 8.2 |
| | 82 - 103 | GQPKRITLASKSSRRKNKKKAK | 4.5 |
| | 82 - 110 | GQPKRITLASKSSRKKKKAKRKQKNHTAE | 3.5 |
| | 103 - 131 | KRKQKNHTAEAAKDKGSDDDDDDEFDKII | 3.7 |
| | 112 -138 | EAAKDKGSDDDDDDEFDKIIQFKKTDI | 5.1 |
| | 112 - 142 | EAAKDKGSDDDDDDEFDKIIQQFKKTDIILK | 4.5 |
| | 241 - 270 | IQRLKRLIRNWGGKDRHLAPNGPGMPHQHL | 3.1 |
| | 243 - 274 | RLKRLIRNWGGKDRHLAPNGPGMPQHLKFTKI | 5.6 |
| | 246 - 278 | RLIRNWGGKDRHLAPNGPGMPQHLKFTKIRDDK | 3.5 |
| | 520 - 554 | RNALLKAFKHHPQLSELFKEKLLGDHALTKDLSID | 4.3 |

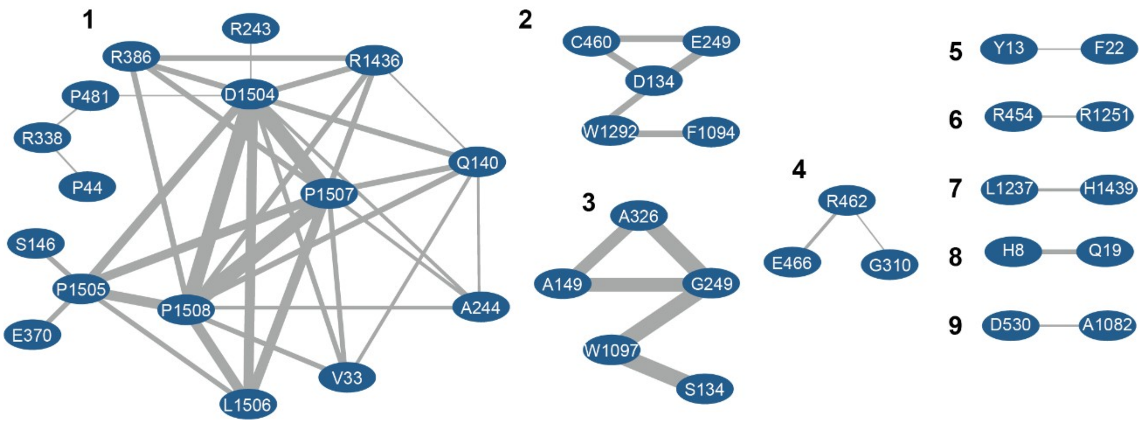


**Figure S1: Community decomposition of the residue coevolution network generated for Tcf25 domain of Rqc1p homologs using PFStats.** Network decomposition identified nine distinct communities, numbered from 1 to 9. Residues numbers correspond to amino acid positions in the global multiple sequence alignment. Edge thickness is proportional to the strength of the coevolutionary association between residues.

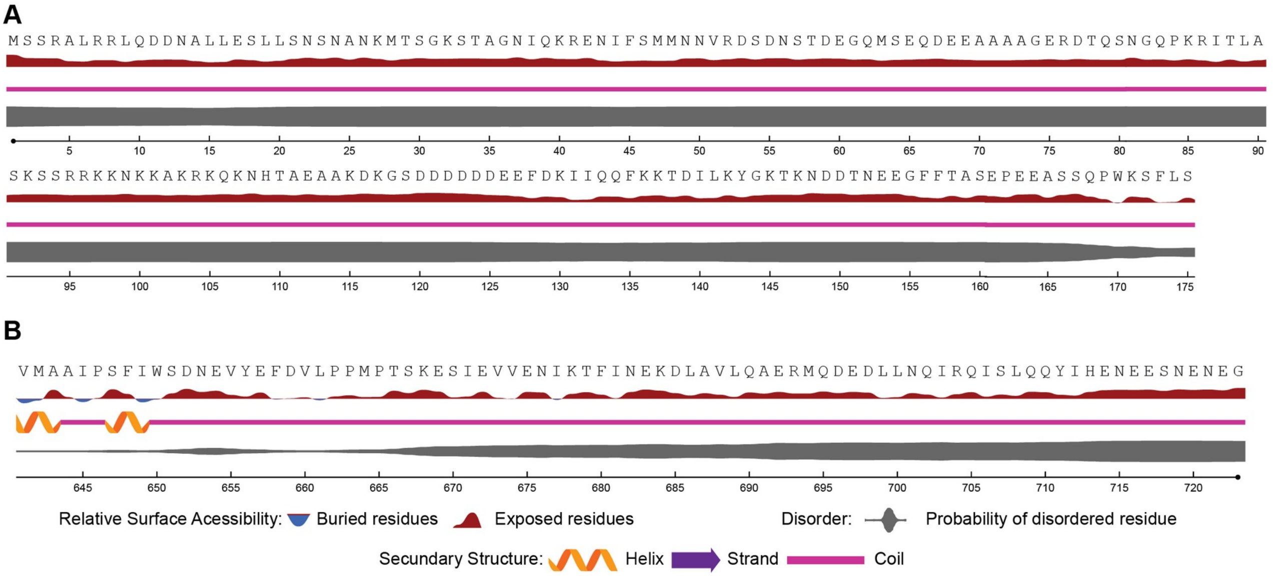


**Figure S2: Secondary structure prediction of Rqc1p reveals disordered regions in N and C-terminal regions.** Predicted secondary structure and structural properties of the N-terminal **(Panel A)** and C-terminal **(Panel B)** regions of Rqc1p obtained using NetSurfP-3.0. Secondary structures are indicated as α-helices (orange), β-strands (purple) and coils (pink). Relative surface accessibility is shown with exposed residues (>0.25, red) and buried residues (<0.25, blue). Disordered regions are represented with line thickness proportional to the disorder probability (gray).

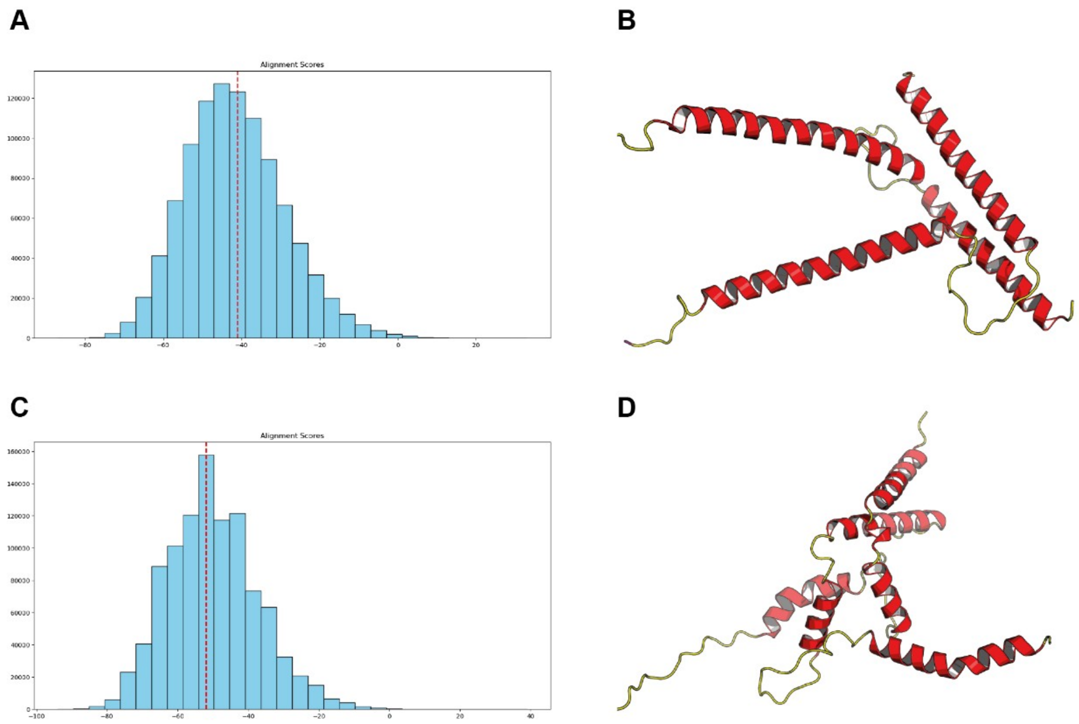


**Figure S3: Negative controls for the putative HLH fold. Panel A:** Alignment significance test using MITF HLH and the Rqc1p N-terminal. The histogram shows the scores for the best alignments between the two segments when one of them is shuffled (repeated $10^3$ times), and the dotted red line corresponds to the best alignment score between MITF HLH and the N-terminal from Rqc1p. **Panel B:** The resulting structure after the Rqc1p N-terminal is modelled against MITF HLH and subjected to energy minimization and molecular dynamics (1ns NVT equilibration and 100ns in NPT conditions). Panels C and D: Same as A and B, using the C-terminal from Rqc1p as a decoy.

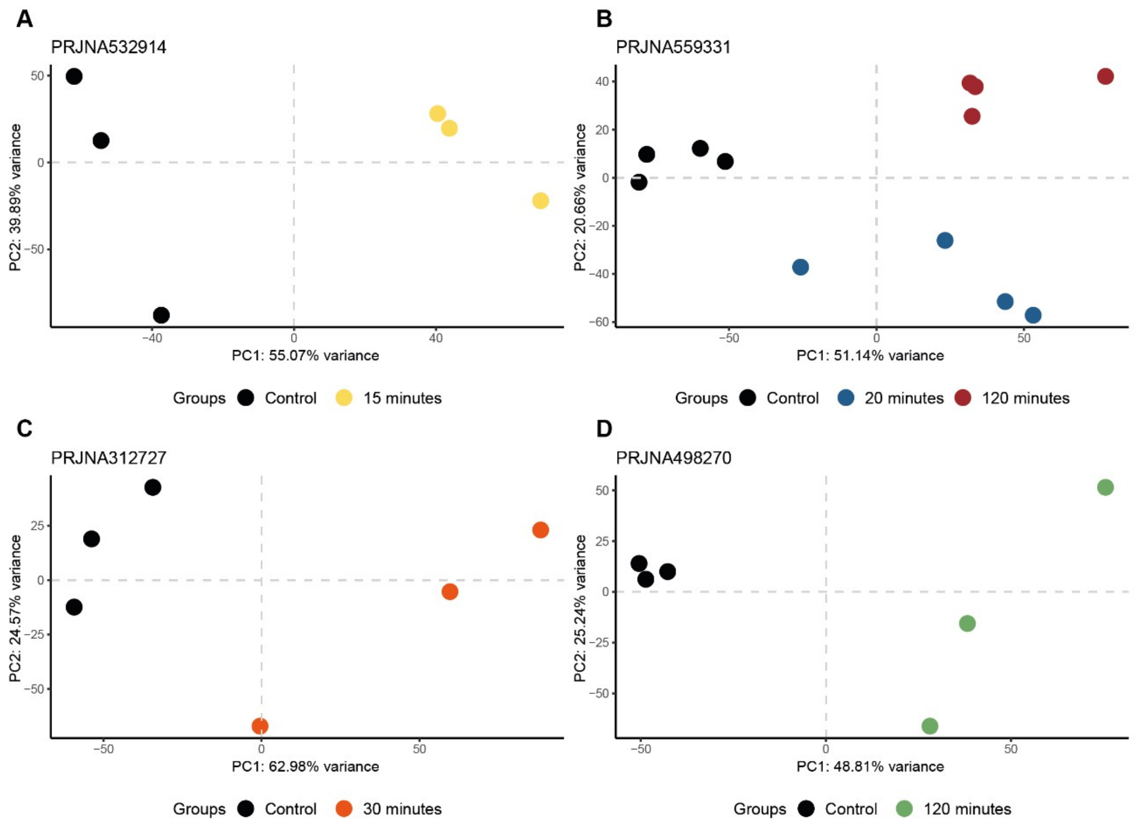


**Figure S4: Principal component analysis (PCA).** Scatter plots showing the PCA results for each study analyzed. Each point represents a sample from the experimental conditions. The principal components PC1 and PC2 explained, respectively: **Panel A:** 55.07% and 39.89% for PRJNA532914; **Panel B:** 51.14% and 20.66% for PRJNA559331; **Panel C:** 62.98% and 24.57% for PRJNA312727; **Panel D:** and 48.81% and 25.24% for PRJNA498270.

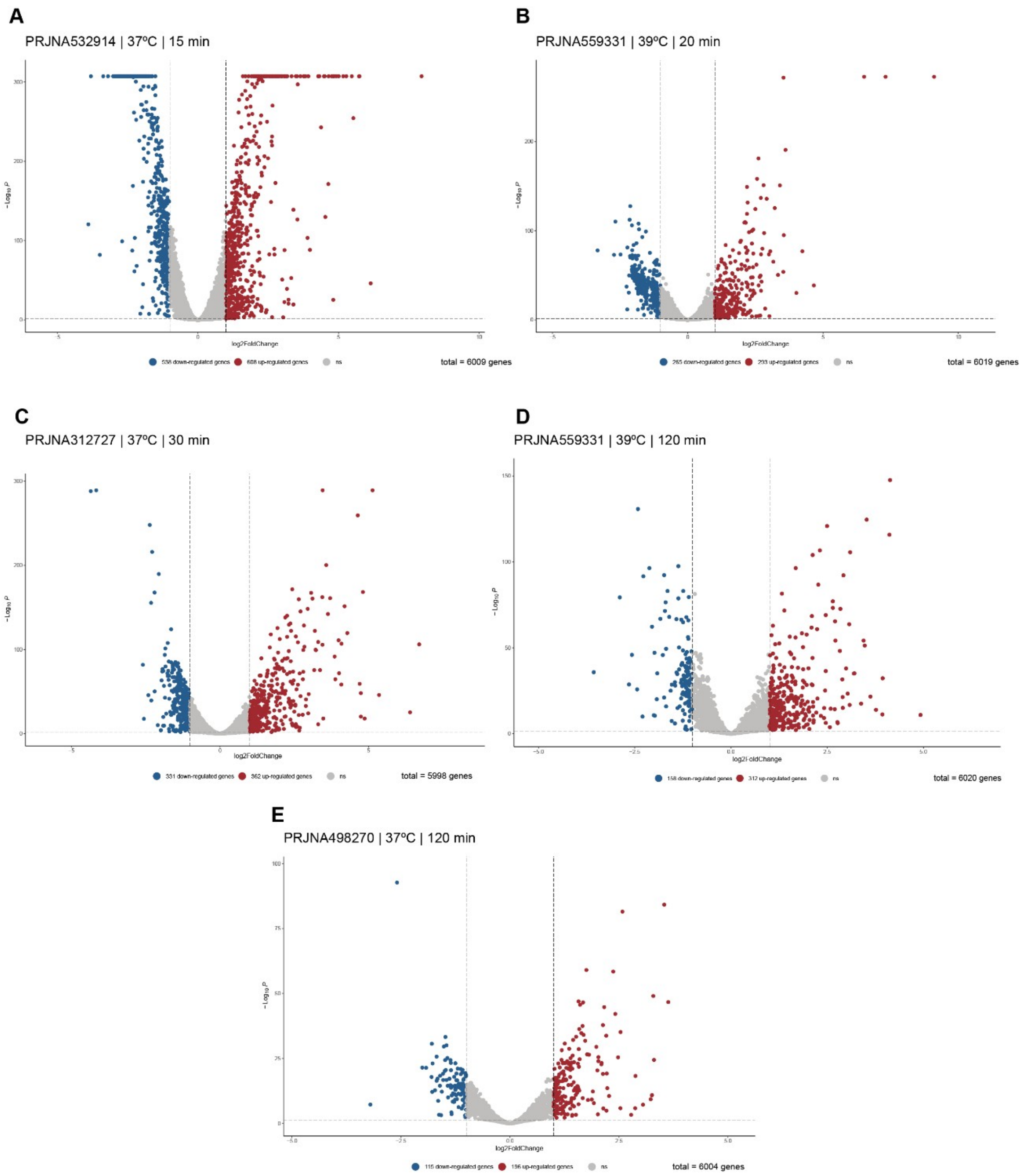


**Figure S5: Volcano plots representing the differentially expressed genes for each study analyzed.** Dots represent non-differentially expressed genes (gray), downregulated genes (blue) and upregulated genes (red) according to log2FoldChange ≥ |1| and padj < 0.05. The studies are ordered according to time in heat shock conditions. **Panel A:** PRJNA532914, 15 minutes at 37°C. **Panel B:** PRJNA559331, 20 minutes at 39°C. **Panel C:** PRJNA312727, 30 minutes at 37°C. **Panel D:** PRJNA559331, 120 minutes at 39°C. **Panel E:** PRJNA498270, 120 minutes at 37°C.
